# Miniscope-LFOV: A large field of view, single cell resolution, miniature microscope for wired and wire-free imaging of neural dynamics in freely behaving animals

**DOI:** 10.1101/2021.11.21.469394

**Authors:** Changliang Guo, Garrett J. Blair, Megha Sehgal, Federico N. Sangiuliano Jimka, Arash Bellafard, Alcino J. Silva, Peyman Golshani, Michele A. Basso, H. Tad Blair, Daniel Aharoni

**Affiliations:** David Geffen School of Medicine, University of California Los Angeles, Los Angeles, CA, 90095 USA; Department of Neurology, David Geffen School of Medicine, University of California, Los Angeles, Los Angeles, CA, USA; Department of Psychology, UCLA, Los Angeles, CA 90095-1563, USA; Integrative Center for Learning and Memory, University of California, Los Angeles, Los Angeles, CA, USA; Department of Psychiatry and Biobehavioral Sciences, University of California, Los Angeles, CA 90095; Department of Neurobiology, University of California, Los Angeles, CA 90095; Jane and Terry Semel Institute for Neuroscience and Human Behavior, University of California, Los Angeles, CA 90095 USA; Brain Research Institute, University of California, Los Angeles, CA 90095; Department of Physiology, David Geffen School of Medicine, University of California, Los Angeles, Los Angeles, CA, USA; West LA Veterans Affairs Medical Center, Los Angeles, CA, USA; Intellectual and Developmental Disabilities Research Center, University of California, Los Angeles, Los Angeles, CA, USA

**Author notes:** These authors contributed equally (Changliang Guo, Garrett J. Blair). e-mail correspondence to Daniel Aharoni.

## Abstract

Imaging large-population, single-cell fluorescent dynamics in freely behaving animals larger than mice remains a key endeavor of neuroscience. We present a large field of view open-source miniature microscope (MiniLFOV) designed for large-scale (3.6 × 2.7 mm), single cell resolution neural imaging in freely behaving rats. It has an electrically adjustable working distance of up to 3.5 mm±100 μm, incorporates an absolute head-orientation sensor, and weighs only 13.9 grams. The MiniLFOV is capable of both deep brain and cortical imaging and has been validated in freely behaving rats by simultaneously imaging >1000 GCaMP7s expressing neurons in the hippocampal CA1 layer and in head-fixed mice by simultaneously imaging ~2000 neurons in the dorsal cortex through a cranial window. The MiniLFOV also supports optional wire-free operation using a novel, wire-free data acquisition expansion board. We expect this new open-source implementation of the UCLA Miniscope platform will enable researchers to address novel hypotheses concerning brain function in freely behaving animals.

---

Understanding how populations of neurons and underlying circuits give rise to complex behavior remains a key endeavor of systems neuroscience. Novel imaging tools combined with the development of new calcium (Ca^2+^) indicators have advanced neuroscience by introducing the ability to monitor large populations of neurons simultaneously and track identified neurons across weeks to months^1^. Decades of research has led to the development and refinement of optical techniques, such as multi-photon^2–4^, confocal^5^, and light sheet microscopy^6^ to image the structures and functions of large neuronal networks with cellular and subcellular resolutions^7^. However, many of these approaches require bulky imaging devices that can only be used on head restrained animals^8,9^, constraining experiments from being conducted in more naturalistic environments and behaviors.

Single photon epifluorescence miniature microscopy^10,11^ and multi-photon miniature microscopy^12–14^ circumvent the constraint of head stabilization while still achieving single cell resolution, enabling these optical techniques in freely behaving animals and expanding the repertoire of behavioral assays that can be used in conjunction with neural imaging. Open-source head-mounted miniaturized epifluorescence microscopes, such as the open-source UCLA Miniscope^15–17^, FinchScope^18^, miniScope^19^, NiNscope^20^, CHEndoscope^21^, MiniLFM^22^, Miniscope3D^23^, and counterpart miniscopes^24^,^25^, have shown that miniaturized microscopes are light enough to be mounted on the head of a rodent and record behaviorally relevant neural signals extracted by further analytical techniques^26–28^. These developments over the past decade have been used extensively in freely behaving animals to reveal neural dynamics related to learning and memory^15^, neurological disorders^17^, and social interactions^29^.

Previous designs of UCLA Miniscopes were developed specifically for mice, which resulted in a field of view (FOV) of ~ 1 mm^2^, limiting their capabilities and applications in imaging large-scale, single cell fluorescent dynamics in larger animals^30^. Furthermore, Ca^2+^ indicators generally have lower expression and fluorescence in animals other than mice, motivating the need for a high-sensitivity, large FOV imaging system with single cell resolution. In addition, current wired miniature microscopy devices lack optional configurations for wire-free operation, impeding experiments that involve recording multiple interacting animals simultaneously or animals navigating large environments^29,31–33^.

Here we report a head-mounted, open-source Large Field of View Miniscope (MiniLFOV) developed as a new implementation of the UCLA Miniscope platform. It has two optical configurations optimized at different working distances (WD) for superficial and deep neural imaging. A 1.8-mm-WD configuration provides 2.5 μm (center) to 4.4 μm (edge) resolution across 3.1 × 2.3 mm FOV, and a 3.5-mm-WD enables 3.5 μm (center) to 6.2 μm (edge) resolution across 3.6 × 2.7 mm FOV. With a high sensitivity 5MP monochrome complementary metal-oxide-semiconductor (CMOS) image sensor, the MiniLFOV yields 20-fold better sensitivity than the previous generation Miniscope V3, and twice the sensitivity compared to current generation Miniscope V4. The WD is electrically adjustable to cover 1.8 mm/3.5 mm±100 μm depth focus using an electrowetting lens (EWL). Incorporated with a 9-axis absolute-orientation sensor, the MiniLFOV has the ability to collect head movement data at up to 100 Hz, which can help investigate head orientation related neural mechanisms^34^.

The MiniLFOV is 35 mm tall, weighs 13.9 grams, and can be modified for different experiments by swapping its custom objective module. The two WD configurations enable multiple applications such as cortical imaging through a cranial window and deep brain imaging using one or multiple implanted optical probes. Furthermore, we also developed a small, wearable wire-free data acquisition (DAQ) expansion board (wire-free DAQ, 3.5 g) which adds wire-free capabilities to all generations of coaxial cabled UCLA Miniscopes. The wire-free DAQ is battery powered by an on-board, single-cell lithium-polymer battery and records imaging data onto an on-board microSD card. The wire-free DAQ can be mounted directly onto the MiniLFOV’s housing or be worn as a backpack. Power, bi-directional configuration commands, and high bandwidth uni-directional imaging data are transmitted between the MiniLFOV and wire-free DAQ via a short coaxial cable. Integrated with an IR remote control receiver, the system supports remote triggering with an IR remote control transmitter.

We validated the wired MiniLFOV in freely behaving rats running on a large rectangular track by imaging GCaMP7s expressing neurons in the hippocampal CA1 pyramidal layer using a 1.8 mm-diameter relay Gradient Refractive Index (GRIN) lens (FOV=~2.54 mm^2^)^22,35,36^, with over 1,300 cells imaged in a single 15-minute session. We also validated the wire-free configuration of the MiniLFOV in rats exploring an open circular field. In addition, we validated the MiniLFOV in head-fixed mice on a circular treadmill by imaging GCaMP6f expressing neurons in dorsal cortex through a cranial window with over 1,900 neurons recorded in a single 7-minute session. With greater than a 30-fold increase in FOV than the previous generation UCLA Miniscope V3 and 12-fold increase in FOV than the current generation UCLA Miniscope V4 (**Video 1**), we expect this imaging platform will generate qualitatively and quantitatively new data for neuroscience applications in freely behaving large animals. As larger animals, rats for example, can often perform more sophisticated behaviors than mice^37^, increasing the yield of neurons recorded simultaneously in these animals has the potential to yield novel discoveries about brain function, particularly when activity is encoded within a subset of the population^38–40^. Importantly, the MiniLFOV is fully open-source, building off the already widely adopted open-source UCLA Miniscope ecosystem, and has been developed to reduce the technical and economic hurdles common in novel tool adoption, making it broadly accessible to the global neuroscience community.

## Results

### System design

The MiniLFOV (**Fig. 1a**) consists of three modules, an objective module, an emission module, and a sensor module, holding the optical components and custom Rigid-Flex printed circuit board (PCB). The PCB consists of an excitation LED sub-circuit, an electrowetting lens tuning and head orientation sub-circuit, a CMOS image sensor sub-circuit, and a power-over-coax and serializer sub-circuit, all interconnected by an embedded flex printed circuit (**Fig. 1a, b**). The 1.8-mm-WD objective module (**Fig. 1c**) contains 3 achromatic lenses (#45-345, #49-656, #45-092, Edmund Optics) and a 3D printed spacer. The emission module (**Fig. 1d**) holds the excitation filter (ET470/40x, Chroma), dichroic mirror (T500spxr, Chroma), emission filter (ET525/50m, Chroma), aspherical lens (#49-658, Edmund Optics), 3D printed lens holder, and concave lens (#45-019, Edmund Optics). The sensor module is designed for holding the EWL driven by an EWL driver (MAX14515, Maxim) and a 5MP monochrome CMOS image sensor (MT9P031, Onsemi), see **Figure 1b**. The main modules are screwed together with M1 thread-forming screws and 0-80 socket head screws.

**Figure 1.**
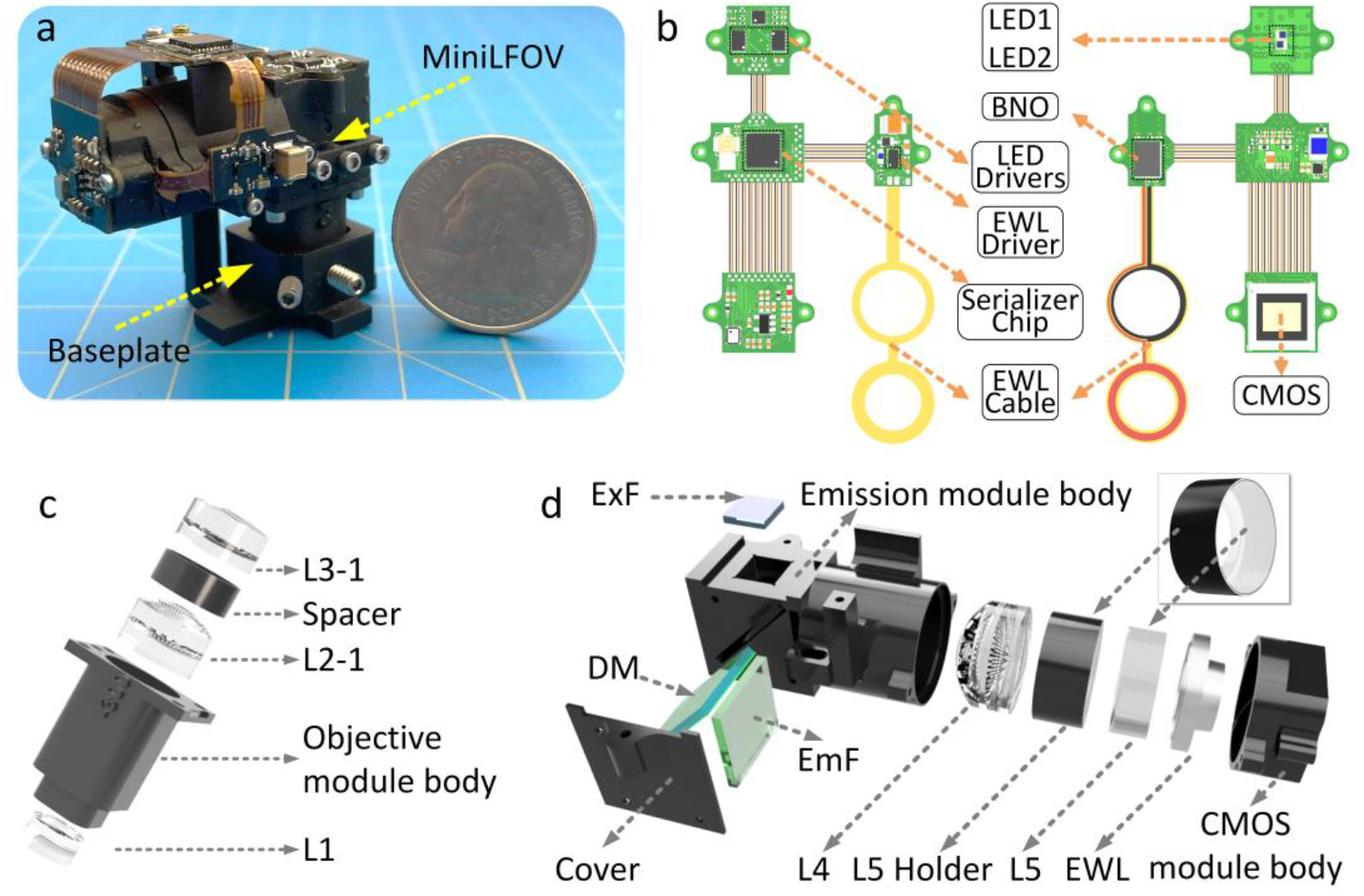
Design of MiniLFOV. (**a**) Photograph of MiniLFOV with baseplate. **(b)** Rigid-Flex PCB of the MiniLFOV consisting of four rigid PCBs connected by an internal flex printed circuit. Two LEDs are housed on the LED circuit board and driven by two led drivers with an I^2^C digital potentiometer for brightness control. An EWL driver and EWL cable holding the EWL is housed along side an absolute-orientation sensor (BNO055) for collecting head orientation data. A 5MP monochromatic CMOS image sensor (MT9P031) is used for capturing Ca^2+^ fluorescence and sending digitized image data to the serializer system. A serializer chip serializes the imaging data and sends it over a coaxial connector to communicate with a custom MiniscopeDAQ system. (**c**) Objective module of the MiniLFOV (1.8-mm-WD configuration) containing a 3D printed objective module body, three achromatic lenses (L1, L2-1, and L3-1) and a spacer between L2-1 and L3-1. **(d)** Emission and sensor module of the MiniLFOV. The emission module consists of a 3D printed emission module body with a 3D printed cover, aspheric lens L4, a plane-concave lens L5, L5 holder, excitation filter (ExF), dichroic mirror (DM), and emission filter (EmF). The sensor module consists of a 3D CMOS module body holding the Electrowetting lens (EWL) and a mounting pocket for the CMOS image sensor.

Power, communication, and image data are packed into a single, flexible 50 Ω coaxial cable (CW2040-3650SR, Cooner Wire) using power-over-coax filtering and a serializer/deserializer pair for bi-directional control communication and uni-direction high bandwidth data streaming (TI DS90UB913A/DS90UB914A).

The MiniLFOV interfaces with the open-source Miniscope DAQ Software to stream, visualize, and record neural dynamics and behavioral data. The software enables multiple Miniscope and behavioral camera video streams, allowing for multi-animal neural and behavioral recordings. This DAQ platform allows for adjustment of excitation intensity, EWL focus, image sensor gain, and frame rate. The DAQ supports synchronizing with external devices through an outputted frame synchronization signal or an input trigger to externally trigger the recording. The list of the components used are given in **Table 3**.

MiniLFOV accessories also include a reinforced baseplate attached to the skull with dental/bone cement which allows consistent mounting of the MiniLFOV, and a protective cap to cover the GRIN lenses implanted. Details of assembling the MiniLFOV are given in **Methods section** and **Figure s1**.

### Optical performance

The optics of the system were designed and optimized using Zemax OpticsStudio. The excitation path (blue in **Fig. 2a**) and emission path (green in **Fig. 2a**) are split by the dichroic mirror, which also folds the emission path to lower the center of mass of the system (**Fig. 2a, f** and **g**). The emission filter is placed in the emission path to block possible excitation light leakage and back-scatter. The high resolution of the system and large FOV are achieved by using 5 off-the-shelf lenses with optimized distance and sequence between them. The 1.8-mm-WD configuration of the system is designed to have a 0.25 NA at the object space, with the Modulation Transfer Function (MTF) curves reaching zero at 500 lps/mm (2 μm) on the image plane The field curvature of the optics in objective space across a 3 mm FOV is 130 μm and the magnification of the system is 1.9 (**Fig. 2f**). The optical system also enables an electrically adjustable WD of ±100 μm in the object space with the EWL (**Fig. 1d, Fig. s2, s3**). To collect the large FOV at its highest resolution, a CMOS image sensor is used with 2592 × 1944 pixels of 2.2 μm pixel size, with a 3.1 mm × 2.3 mm FOV achieved (**Fig. 2b, s2**). In practice, the achieved spatial resolution of the system is 2.5 μm (**Fig. 2b, c**) at the center of the FOV, dropping to 4.4 μm at the edge. The resolution is also validated with full width at half maximum (FWHM) values of 1-μm beads imaged (**Fig. 2d, e, s3, s5**). Driven with a 66.667 MHz pixel clock, the sensor runs at 11 frames per second (FPS) at full resolution and 23 FPS with 2× horizontal and vertical binning. With 2× pixel binning, 4.4 μm resolution is achieved at the center of the FOV (dropping to 6.2 μm at the edge) and is the configuration used for all data collection shown in this paper. For higher FPS (up to 15 FPS at full resolution and 30 FPS at 2× binned resolution), a 96 MHz clock can be used but requires shorter or larger diameter coaxial cables for stable use. Higher FPS can additionally achieved by cropping the recording window or increasing the binning factor. To optically excite Ca^2+^ indicators across its large FOV, two LEDs (LXZ1-PB01, LUXEON) are available in the excitation path with a band-pass excitation filter placed below to narrow their spectral band. With 2 × 2 pixel binning and maximum analog gain (MiniLFOV_bin2×; gain=8×), the MiniLFOV has twice the sensitivity than that of a Miniscope V4 at maximum analog gain (gain=3.5×) resulting in a 2× decrease in excitation power needed per unit area. A comparison of the sensitivity and signal-to-noise ratio (SNR) of Miniscope V3, Miniscope V4, and MiniLFOV is shown in **Figure s4**. We have also validated that there is no difference in SNR for regions of interest in the center of FOV and the periphery testing with Negative USAF 1951 Hi-Resolution Target without GRIN lens inserted between the MiniLFOV and the Target.

**Figure 2.**
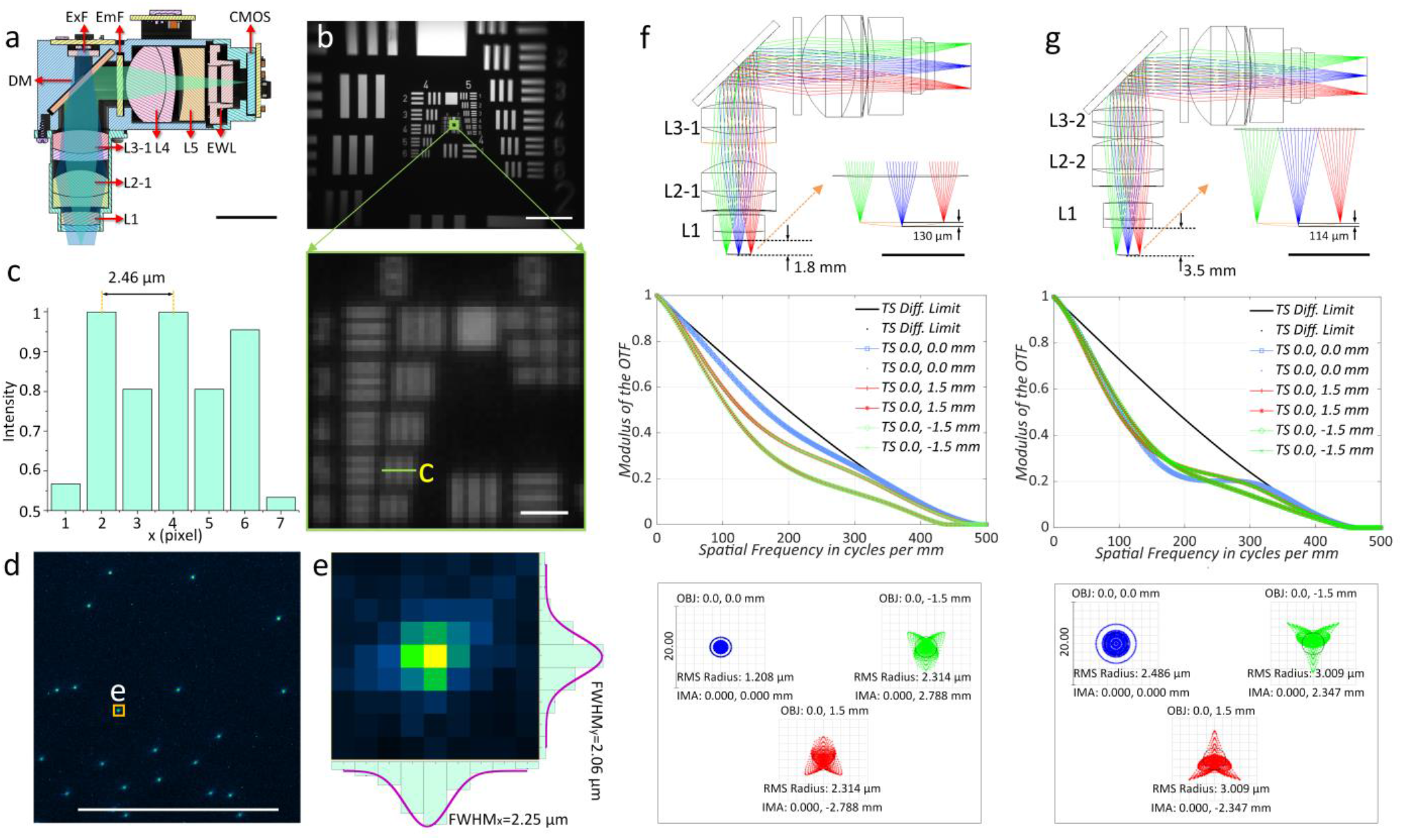
Optical design and performance of the MiniLFOV. (**a**) Cross section of the MiniLFOV assembly, showing the optical components inside the body, including objective module (1.8-mm WD) with 3 achromatic lenses (L1, L2-1, and L3-1), emission module with 1 achromatic lens (L4), 1 concave lens (L5), and 1 electrowetting lens (EWL). It also includes excitation filter (ExF), Dichroic mirror (DM), and emission filter (EmF). Excitation path from 2 LEDs is highlighted in blue and emission path is highlighted in green. Scale bar: 10 mm. (**b**) The 3.1 × 2.3 mm FOV is shown using a Negative USAF 1951 Hi-Resolution Target (#55-622, Edmund Optics). Scale bar: 500 μm (**top**), and 10 μm (**bottom**). (**c**) 2.5 μm (Group 8 Element 5, green box; 406 lps/mm) can be resolved experimentally from the cross-sectional profile along the green line in (**b, bottom**). (**d**) 1.8-mm-WD objective module configuration is shown for use in deep braining imaging. (**d**) 1-μm fluorescent beads imaged with the MiniLFOV. Scale bar: 250 μm. (**e**) xy sections of one selected bead marked in **d** with FWHM_*x*_ =2.25 μm and FWHM_*y*_=2.06 μm respectively. (c,d) Normalized and averaged lateral(**f**) Zemax simulation of emission path shows the 1.85 mm-WD-configuration with 130 μm field cuvature in the object space (**top**), MTF of the three positions on the image plane (**middle**), and spot diagram of the optics (**bottom**). The values of this function reach zero when x gets to 500 lps/mm, i.e. 2 μm resolution, at the image plane (1.1 μm in the object space), which closely agrees with measured results of the system. As shown in the spot diagram of the optics at the image plane, RMS radius is 1.208 μm at the center and 2.134 μm at (0, 1.5 mm/−1.5 mm). The magnification of the optics is given by 2.788/1.5=1.86 calculated from the spot diagram, see **(e, bottom)**, in which the points (0, 1.5 mm/−1.5 mm) are focused at (0, 2.788 mm/-2.788 mm) on the CMOS sensor. (**g**) Zemax simulation of emission path shows the 3.5-mm-WD configuration with 114 μm field curvature in the object space (**top**). MTF curves reach zero when x axis gets 450 lps/mm, i.e. 2.2 μm at the image plane (1.4 μm in the object space) (**middle**). 3.5-mm WD (right) has RMS radius 2.486 μm at the center and 3.009 μm at (0, −1.5 mm/1.5 mm). The magnification is calculated by 2.347/1.5=1.56 (**bottom**), with 3.6 mm × 2.7 mm FOV achivable. experimentally.

The 3.5-mm-WD configuration (**Fig. 2g**) uses L1, L2-2 (#49-657, Edmund Optics), and L3-2 (#45264, Edmund Optics) in the objective module and is designed to give additional space below the MiniLFOV for imaging through cranial windows or other thick samples. MTF curves reach zero at 450 lps/mm (2.2 μm) at the image plane, and the RMS radius is 2.5 μm on-axis (x, y = 0 mm) and 3.0 μm off-axis (x=0, y=1.5 mm; x=0, y=−1.5 mm). In practice, the achievable resolution is 3.5 μm, or 362 lps/mm, at the center of the FOV with full resolution at 11 FPS. A resolution of 6.2 μm is achievable with 2x horizontal and vertical binning to run at an acquisition rate of 23 FPS. The magnification is 1.56, with which results in a full FOV reaching 3.6 × 2.7 mm.

### Imaging place cells in hippocampal CA1

Studies of the neural representations of space have been prominent ever since the discovery of place cells in the 1970’s^41^. Many pyramidal neurons within the dorsal hippocampus of rats respond when the animal occupies a particular location within an environment^42^. Neurons with such a spatial response are called “place cells” and it is believed the hippocampus constructs an internal map of an environment (a “cognitive map”) which is crucial for spatial navigation and memory^11^. Due to the laminar nature of the CA1 pyramidal layer and relative ease of access using implanted optical probes, many experiments on spatial navigation and episodic memories have begun using miniaturized endoscopy in this brain region^11,15^. However, most experiments have been restricted to mice. Here we performed Ca^2+^ imaging in the hippocampal dorsal CA1 region of rats to demonstrate the feasibility and extended capabilities of MiniLFOV recordings in freely behaving larger animal models.

Currently available open-source UCLA Miniscopes support imaging up to around a 1 mm^2^ FOV, which is sufficient to yield hundreds of imaged neurons in the mouse hippocampus^17^. However, in animal models that are larger than mice, a larger FOV is often needed to achieve similar cells counts. Furthermore, Ca^2+^ imaging in larger animals can pose additional difficulties: larger brain sizes can produce greater tissue scattering, and GCaMP expression can be suboptimal using commercially available viral vectors that are often optimized for use in mice. A high sensitivity, large FOV miniscope system is needed to ameliorate these problems.

To record place cell activity from the dorsal CA1 region in rats, we imaged Ca^2+^ dynamics in GCaMP7s-expressing neurons in dorsal CA1 (**Fig. 3a, s6, s11**) while rats performed navigation of a rectangular track environment. An overview of installing and removing the MiniLFOV is shown in **Figure s6–s7**. At around a total mass of 20 grams (which includes the MiniLFOV, baseplate, and cement), rats can easily wear the whole device without any signs of difficulty when running (**Fig. 3b**). In total, imaging data from 15 rats across months of recordings were recorded with the MiniLFOV in this task, but we present one exemplary session of one rat to demonstrate the effectiveness of the system. Example histology from 15 rats, showing GCaMP expression with the pyramidal layer for CA1 beneath the implanted lens location is in **Figure s8**. The average running speed along the center of the maze and their reward rate (rewards/min) in the session is presented to show the MiniLFOV doesn’t restrict the freely moving of the mice (**Figure. s9**). **Figure 3c** shows a maximum projection image from an example 15-minute motion corrected video recording captured by the MiniLFOV. The raw video is recorded at 23 fps and around 15 minutes with 972 × 1296 pixels for each frame after within-sensor 2x pixel binning and then further cropped to the pixel region containing the relay GRIN lens (720 × 720 pixels). In this example recording, a total number of 1357 cells can be detected and extracted using extended constrained nonnegative matrix factorization (CNMF-E) analysis via Calcium Imaging data Analysis (CaImAn)^27^,^28^ (**Fig. 3d**). Recordings took place on a rectangular track (2.5 m × 1.25 m; **Fig. s10**), with two reward feeders on two corners of the track. Positions on the track (**Fig. 3e**) and head orientation (**Fig. 3f**) of the rat were extracted from a software synchronized behavioral camera and on-board head orientation sensor respectively (see **Methods** and **Fig. s10)**. Low-pass filtered Ca^2+^ traces of 25 example cells are shown in **Figure 3g**.

**Figure 3.**
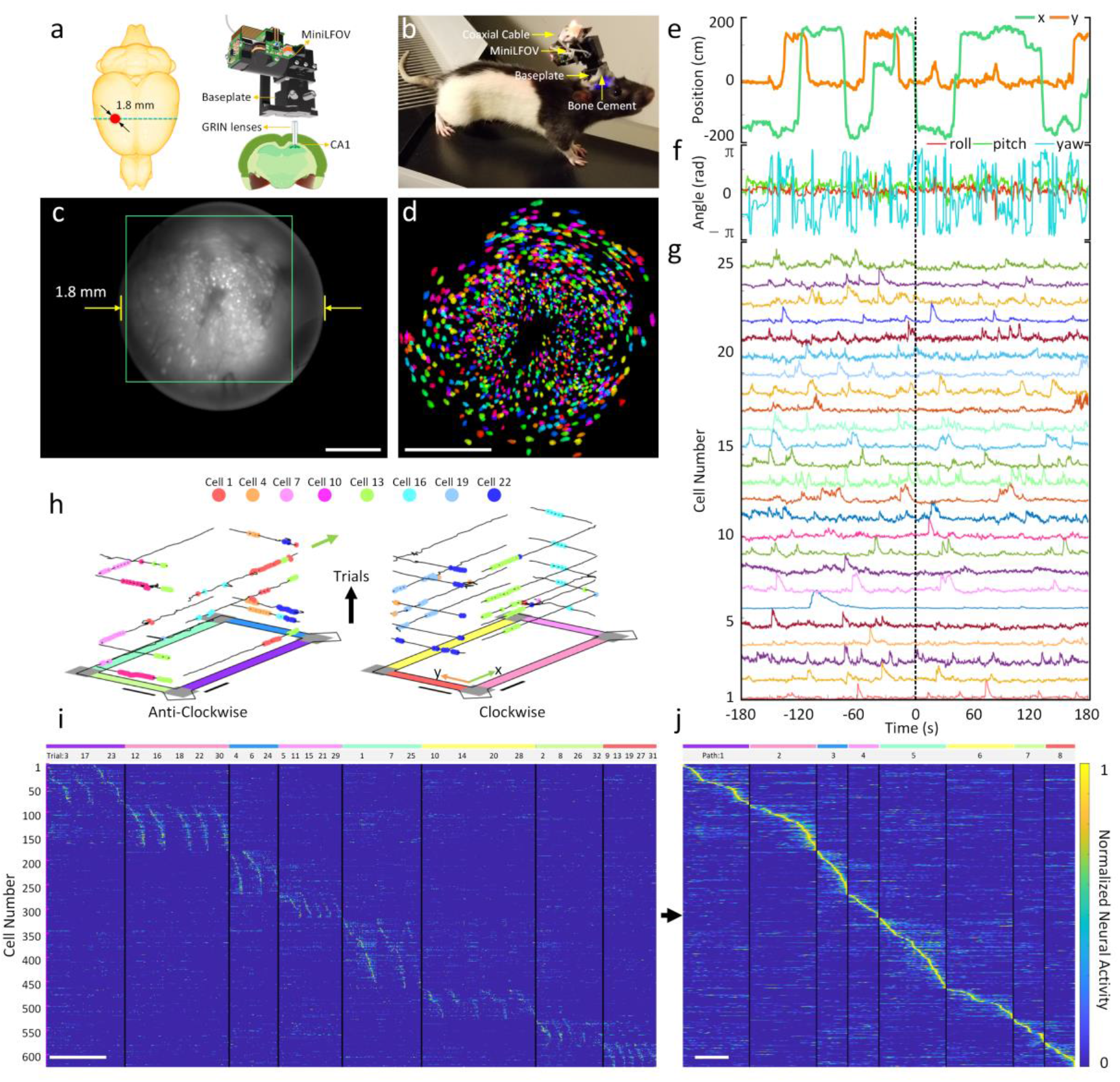
Strategy for imaging Hippcampal dorsal CA1 in freely behaving rats. **(a)** Schematic of the imaging area in the rat brain. A custom relay GRIN lens is implanted above CA1 to relay fluorescence to the intermediate image plane formed between the top of the GRIN lens and bottom of the MiniLFOV. **(b)** Rat with MiniLFOV mounted on the skull via the reinforced MiniLFOV baseplate (**c**) Maximum projection from a 15-minute single recording session after motion correction. Scale bar: 500 μm. (**d**) Pixel area outlined in the green box in **c** showing extracted contours of cells (N=1357) from CNMF-E analysis via CaImAn, colored randomly. Scale bar: 500 μm. **(e)**Example x and y track position as the rat runs in the 2D environment. (**f**) Example head orientation in terms of the roll, pitch, and yaw Eular angles. **(g)** Example extracted Ca^2+^ traces with CaImAn pipeline for 25 cells recorded during the session after low-pass filtering. **(h)** Deconvolved neural activity of 8 example place cells spatially plotted on top of traversals along the track divided up by trail number and direction. Scale bar: 50 cm. **(i)** 626 place cells’ normalized deconvolved neural activity across all arm traversals sorted by the location of peak spatial activity rate shown in **j**. Data shown here is speed thresholded (20 cm/s) to remove bouts of stationary active along traversals. Scale bar: 10 s. **(j)** Normalized spatial neural activity rate along all paths for these 626 place cells. Scale bar: 125 cm.

As expected, a significant population of the cells (626; 46.2%) are spatially tuned along at least one direction on the 4 arms on the rectangular track. A cell is considered a place cell if it has significantly higher spatial information than chance (spatial information above at least 95% of 500 randomly shifted deconvolved neural activity for each cell)^43^. The behavior of the rat was serialized into 8 paths (4 arms with 2 running directions) (**Fig. 3h-j**). Place cells recorded during the session were significantly modulated by both position and direction of travel and span across all regions of the track (**Fig. 3i-j**). An example video showing the position of the rat, head orientation, and place cell activity during behavior is given in **Video 2 and 3**. These data demonstrate how the MiniLFOV can improve experimental efficacy by yielding more cells in larger research models, broadening the horizon of potential hypotheses for researchers. Demonstration of the MiniLFOV’s capability in terms of consistently recording activity from the same population of cells across multiple sessions is shown in **Figure s11**.

### Optional Wire-free recording in rats

The MiniLFOV can record neural activity without the constraint of being tethered to DAQ hardware mounted off the animal. By mounting a novel wire-free DAQ (3.5 g) system to the side of the MiniLFOV or on a backpack and connecting it to the MiniLFOV through a 4 cm long, 50 Ω coaxial cable any wired UCLA Miniscope can operate in a wire-free configuration (**Fig. 4a, b**). A single-cell 400 mAh lithium-polymer battery (7.5 g) is used to power the wire-free DAQ and MiniLFOV (supports close to 1 hour of continuous recording) and a microSD card (Class 10 UHS-I microSD, Kingston) is used for on-board data and configuration storage (**Fig. 4b-d**). Once the wire-free DAQ is powered on, an on-board microcontroller (MCU) (ATSAME70N21A, Microchip) reads configuration data from the microSD card and then implements that configuration in the MiniLFOV. Configuration parameters include excitation intensity, EWL focus, and frame rate, gain, and FOV window of the image sensor. A status LED is integrated onto the wire-free DAQ for displaying current device state and visually synchronizing recording with behavior cameras (**Fig. 4c**). The wire-free DAQ uses an infrared (IR) remote control receiver to receive digital commands, encoded into a 38 KHz IR carrier frequency, from an IR transmitter to trigger recording remotely. This implementation of IR communication allows for one-way, wireless data transfer to the wire-free DAQ and MiniLFOV. During recording, data from the attached MiniLFOV is streamed into the memory of the MCU by a direct memory access channel. Each acquired frame is timestamped and then saved onto the microSD card. The data acquisition rate of the wire-free DAQ is limited to a pixel clock rate of around 24MHz by the on-board MCU’s parallel capture mode. Additionally, the MCU can maintain a maximum of ~11 MBps write speed to the microSD card, placing additional constrains on the rate of image acquisition and storage. Due to these constrains, we chose the recording window size to be 608 × 608 pixels at 15 FPS (although it can achieve 20 FPS), corresponding to 1.4 × 1.4 mm^2^ FOV in object space (relay GRIN lens is 1.8 mm in diameter). In the configuration used, around 5.5 MB are written to the SD card per second, allowing a 64 GB microSD card to support multi-hour recording.

**Figure 4.**
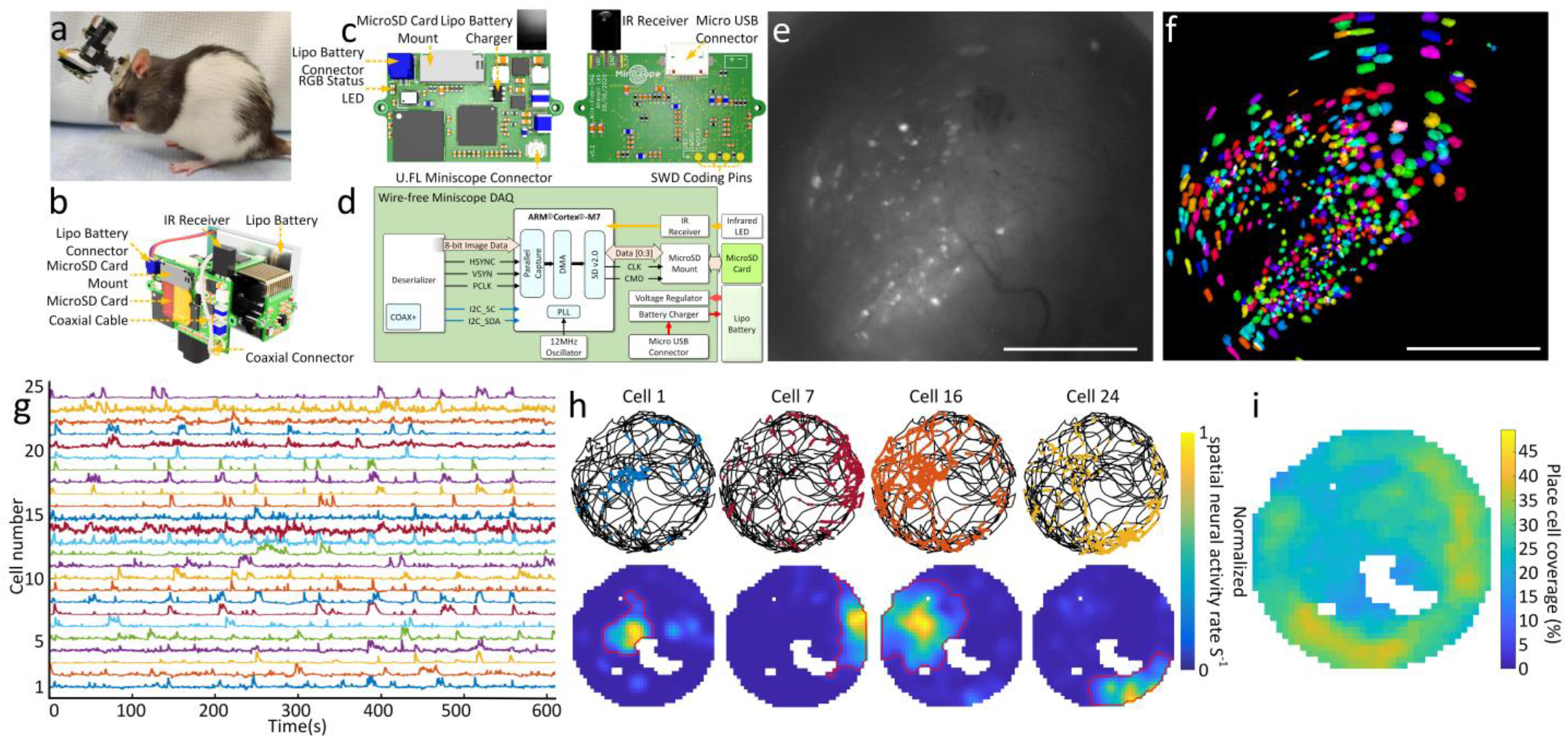
Wire-free imaging of hippocampal CA1 neurons in freely behaving rats in a circular open-field. **(a)** Photograph of a rat wearing MiniLFOV with wire-free DAQ and single-cell lithium-polymer battery. **(b)** 3D rendering of the MiniLFOV equiped with wire-free DAQ and single-cell lithium-polymer battery. **(c)** Top and bottom PCB rendered layouts. **(d)** Block schematic of wire-free DAQ. **(e)** Maximum projection from a 14-minute recording session after motion correction. Scale bar: 500 μm. **(f)** Contours of extracted cells (N=575) from CNMF-E analysis via CaImAn, colored randomly. Scale bar: 500 μm. **(g)** Low-pass filtered Ca^2+^transients from 25 example cells from **f. (h)** Examples of spatially modulated neural activity for 4 place cells during exploration in the open-field. Top: Speed thresholded spatial location of deconvolved neural activity (colored dots) superimposed over rat’s trajectory (black line). Bottom: Binned spatial neural activity rate maps for the same example cells. Red contour denotes the spatial outline of a place field based on a greater than 5% threshold of the binned spatial neural activity rate. **(i)** Combined open-field place field coverage from all 438 detected place cells (76%) showing full coverage of the environment with overall increased coverage near edges.

The feasibility and capability of the wire-free MiniLFOV configuration are validated with an example rat freely moving in a circular open-field (80-cm diameter) as unrestricted behavior and Ca^2+^ dynamics in GCaMP7s-expressing neurons in dorsal CA1 are recorded simultaneously. The maximum projection of a motion corrected 14-minute recording session is shown in **Figure 4e** with a total of 575 extracted cells (**Fig. 4f)**. Low-pass filtered Ca^2+^ traces from 25 example cells are shown in **Figure 4g**. A significant population of cells (N=438; 76%) satisfied our place cell criteria with spatial information above at least 95% of 500 circularly shuffled sessions. Four example place cells show the rat’s trajectory (black lines) and spatial location of deconvolved neural activity (colored dots) (**Fig. 4h (top))** showing clear place preference of neural activity. These example cells’ spatial neural activity rate maps are shown in **Figure 4h**. The red outlined contour denotes the edge of a place field calculated by detecting a 5% cutoff of the binned spatial neural activity rate surrounding the place field. The combined open-field place field coverage from all 438 detected place cells shows full coverage of the circular open-field with overall increased coverage near the edges (**Fig, 4i)**. This wire-free configuration enables untethered behavior during neural imaging in larger animals and has the potential to improve naturalistic behavior by removing cable torque and looming cues. In addition, the wire-free DAQ extends the capabilities of the UCLA Miniscope to simultaneous multi-animal recordings and experiments where tethering is infeasible.

### Imaging cortex in head-fixed mice

With single cell resolution across the entire 3.6 mm × 2.7 mm FOV (3.5-mm-WD configuration, see **Figure. 2g**), the MiniLFOV is capable of large-scale imaging of cortical ensembles in head-fixed mice, freely moving rats, and larger animals. Here, we demonstrate the MiniLFOV enables single cell resolution across the entire FOV (**Fig. 5a**) by imaging GCaMP6f-expressing dorsal cortex^44,45^ through a 4 mm × 4 mm cranial window in head-fixed mice running on a 29-mm circular treadmill (**Fig. 5a, s12a**). A MiniCAM (**Fig. s12a**), an open-source behavior camera developed by the UCLA Miniscope group, was used to record the movement of the mouse (**Fig. s12b**). Maximum-intensity projection image of the raw video after motion correction with background removed is shown in **Figure 5b**. The bright white spots in the image are putative cells. Contours of 1907 extracted cells from CNMF-E analysis via CaImAn circled in yellow distributed across the full FOV. To compare the number of cells Miniscope V4 and MiniLFOV can capture, we randomly chose 500 1-mm diameter circular regions, which is the FOV of Miniscope V4, to calculate the distribution of the cell number. The mean cell number is 211.032 and the standard deviation is 190.567 (**Fig. 5c**). As a comparison, MiniLFOV captured 9 times number of cells on average. To show the MiniLFOV can resolve single cells, 4 sub-regions located near the four corners in the entire FOV are chosen and magnified in **Figure 5d-g**, and background removed to highlight the putative cells (bright white spots in **d-g**) and the contours of the extracted cells (circled in yellow). In each sub-region, 15 cells are randomly chosen with extracted Ca^2+^ traces plotted on the right to show the visual assessment of the SNR during the session after low-pass filtering. Both the contours and the corresponding Ca^2+^ traces have validated the performance of MinilFOV with respect to single-cells resolution. Similar results from a second mouse are shownshown in **Figure s13**.

**Figure 5.**
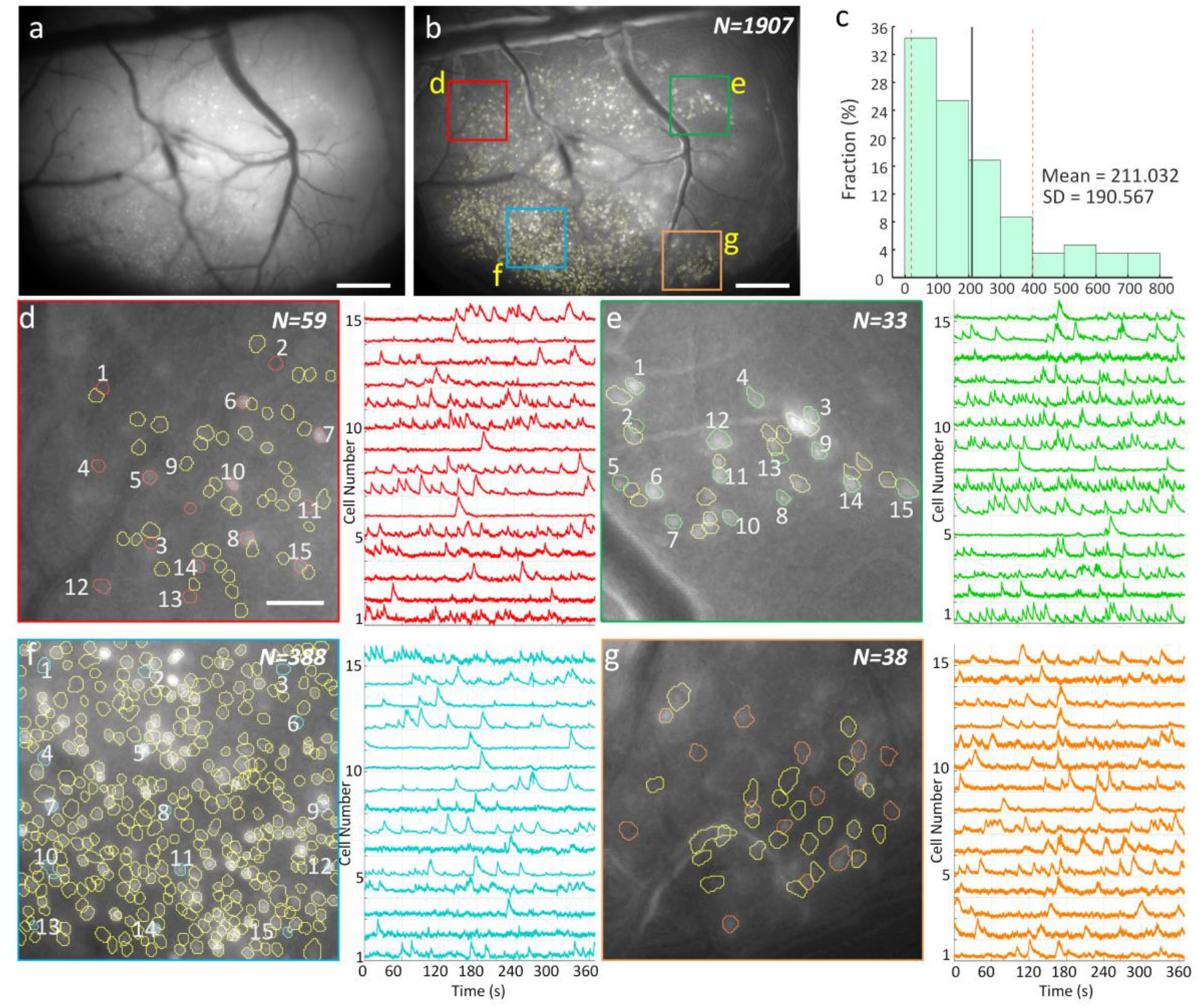
Ca ^2+^ imaging of dorsal cortex through a 4 mm × 4 mm cranial window in head-fixed mice. (**a**) Maximum projection from a 7-minute recording session after motion correction. Scale bar: 500 μm. (**b**) Maximum-intensity projection image of the raw video after motion correction and background removed with contours of 1907 extracted cells from CNMF-E analysis via CaImAn circled in yellow. The colored boxes indicate four sub-regions which are zoomed in and shown in **d-f**. Scale bar: 500 μm. (**c**) Distribution of cell numbers inside 500 randomly chosen 1 -mm diameter circular regions (FOV of Miniscope V4) in the entire FOV of the MiniLFOV. The mean cell number is 211.032 and the standard deviation is 190.567. (**d**) Map of 59 cells in the red-boxed area in **b** and 15 randomly chosen cells with their low-pass filtered Ca^2+^ transients for the numbered cells are shown in red. Scale bar, 100 μm. (**e**) Map of 33 cells in the green-boxed area in **b** and 15 randomly chosen cells with their low-pass filtered Ca^2+^ transients for the numbered cells are shown in green. (**f**) Map of 388 cells in the cyan-boxed area in **b** and 15 randomly chosen cells with their low-pass filtered Ca^2+^ transients for the numbered cells are shown in red. (**g**) Map of 38 cells in the orange-boxed area in **b** and 15 randomly chosen cells with their low-pass filtered Ca^2+^ transients for the numbered cells are shown in cyan.

## Discussion

We present a large field of view (FOV) open-source miniature microscope platform aimed at extending the capabilities of microendoscopic imaging approaches in large rodents and non-human primates, such as rats, marmosets, and macaque monkeys. The system can image up to a 3.6 mm × 2.7 mm FOV at 23 FPS with single cell resolution, has an electrically adjustable WD of up to 3.5 mm±100 μm, incorporates an absolute head-orientation sensor, and weighs under 14 grams, which is under the weight of many head-mounted neural recording devices for rats like micro-drives. The MiniLFOV achieves a 20-fold increase in sensitivity compared to the previous generation Miniscope V3, and a 2-fold increase in sensitivity compared to the current generation mouse Miniscope V4. The MiniLFOV includes a modular objective lens configuration, enabling multiple imaging approaches for deep and superficial brain imaging. The MiniLFOV can be easily attached to and removed from a chronically implanted baseplate that has been designed to provide rigid mechanical support for stable recording in freely behaving animals larger than a mouse. The baseplate mounting mechanism also provides an accurate and repeatable mounting to image the same neural population and track individual neurons across recording sessions (**Figure s6–s7, s11**).

The MiniLFOV has been validated in freely behaving rats by imaging GCaMP7s expressing neurons in the dorsal CA1 layer of the hippocampus (with 1.8-mm GRIN lenses implanted down to 2.5 mm) and is one of first demonstrations of large-scale recordings of place cells in freely behaving rats. Users also have the option of using thinner relay GRIN lenses for deeper brain imaging which are all compatible with the MiniLFOV. Previous recordings of freely behaving rats using original Miniscope V3 designed for mice were severely limiting, typically yielding around 100 cells or fewer per recording^40^. With the presented MiniLFOV, we can now achieve simultaneous recordings of over 1,000 neurons in freely behaving rats during a single 15-minute session (**Fig. 3**). When imaging across the entire FOV of the MiniLFOV, close to 2,000 active neurons could be imaged across the dorsal cortex in head-fixed mice (**Fig. 5**). This platform greatly extends Ca^2+^ imaging capabilities in freely behaving larger animal models and opens new avenues of research that were previously limited by low simultaneous cell counts. Additionally, the entire open-source UCLA Miniscope platform has also been extended for wire-free operation. This has the potential to reduce behavior interference (such as cable tangling and unintentional looming cues) as well as enable recordings from multiple animals socially interacting^29^ and animals exploring large environments. Performance and capabilities of three configurations of the MiniLFOV, i.e., 1.8-mm-WD, 3.5-mm-WD, and wire-free, are listed in **Table 1**. Furthermore, with independently controlled dual-LED excitation, the MiniLFOV can support future two-color excitation configurations by changing one of the LEDs as well as changing filters and dichroic mirror^49^. With a mass of only 13.9 grams, it is feasible for two MiniLFOVs to be mounted onto the skull of a rat for multi-region imaging. The MiniLFOV broadens the application of Ca^2+^ imaging to larger, freely behaving animals than previously realized, enabling large-scale and high cell-count recordings in both cortical and deep brain structures. Moreover, the MiniLFOV, wire-free DAQ, and UCLA Miniscope online resources provide an open-source, accessible, and cost-effective option for neuroscientists seeking to adopt Ca^2+^ imaging into their labs.

## Supporting information

Field-of-view comparison across Miniscope versions

Hippocampal CA1 dynamics in freely behaving rat 1

Hippocampal CA1 dynamics in freely behaving rat 2

## Online Methods

### MiniLFOV design, manufacturing, and assembly

#### Design

The optical system of the MiniLFOV was designed using Zemax OpticsStudio. The goal of the optical system is to produce a miniature microscope with large FOV and high numerical aperture (NA), in which the desired NA is set to be 0.25 in the objective space and FOV to be a minimum of 3 mm. Lenses are chosen to achieve the optical performance while maintaining a compactable and lightweight size. The distance between lenses is optimized to achieve single cell resolution across the entire FOV with minimal field curvature. Three achromatic lenses L1 (d=6.25 mm, focal length=60 mm, #45-345, Edmund Optics), L2-1 (d=9 mm, focal length=12 mm, #49-656, Edmund Optics), and L3-1 (d=9 mm, focal length=27 mm, #45-092, Edmund Optics) are used to build the 1.8-mm-WD objective module with a 3.5 mm tall spacer placed between L2-1 and L3-1. L1 (d=6.25 mm, focal length=60 mm, #45-345, Edmund Optics), L2-2 (d=9 mm, focal length=18 mm, #49-657, Edmund Optics), and L3-2 (d=9 mm, focal length=36 mm, #45264, Edmund Optics) are chosen to build the 3.5-mm-WD objective module (**Fig. 2g**). One aspherical lens (d=12.5 mm, focal length=14 mm, #49-658, Edmund Optics) and a concave lens (d=12 mm, focal length=-48 mm, #45-019, Edmund Optics) with a lens holder are placed into the emission module body to form the tube lens of the MiniLFOV. The EWL (Corning Arctic 58N, Varioptics/Corning) is placed into the sensor module body (**Fig. 1d, Fig. s1**) for electronic focus adjustment. The MiniLFOV module bodies (objective module body, emission module body, sensor module body), filter cover, baseplate, and protective cap are designed in Autodesk Fusion 360 (educational license) and printed with black resin (FLGPBK04, Formlabs) by a table-top stereolithographic (SLA) 3D printer (Form 3, Formlabs) to produce a lightweight, compact, and low-cost assembly. Three filter slots are designed on the emission module body for mounting a custom diced excitation filter (4 × 4 × 1.1 mm, ET470/40×, Chroma), a dichroic filter (14 × 10 × 1 mm, T500spxr, Chroma) and an emission filter (10 × 10 × 1 mm, ET525/50m, Chroma). All the optical components can be easily assembled without the need for any epoxy or optical glue. A list of the optical components used is given in **Table 3**. The circuit schematic and Rigid-Flex PCB layout are designed using KiCad, a free software suite for electronic design automation (EDA). The PCB is divided up into 4 rigid sub-circuits which include an excitation LED circuit, an electrowetting lens tuning and head orientation circuit, a CMOS image sensor circuit, and a power-over-coax and serializer circuit. The modularity of the PCB design enables quick modification or redesign of individual sub-circuits without the need for modifying the entire PCB layout. The four sub-circuits are connected by a double-sided embedded flex printed circuit (**Fig. 1a, b**). The assembled objective module is attached to the emission module body with five 18-8 Stainless Steel Socket Head Screws (92196A052, McMaster-Carr). The emission module, sensor module, cover, and Rigid-Flex PCB are fastened together with M1 thread-forming screws (96817a704, McMaster-Carr) (see **Fig. s1**). The open-source behavior camera (MiniCAM) for behavior tracking is developed by UCLA Miniscope group (https://github.com/Aharoni-Lab/MiniCAM) and compatible with the open-source UCLA Miniscope DAQ hardware and software. The MiniCAM consists of an M12 optical lens mount, a custom printed circuit board housing a CMOS image senor and supporting electronics, LED illumination ring with 16 red LEDs, and a 3D printed case. The brightness of the LEDs can be adjusted through software for the optimal illumination in dark environments. The MiniCAM runs at around 50 FPS with 1024 × 768 resolution and saves the behavioral data in AVI file format using an MJPG compression video codec directly supported within the UCLA Miniscope software.

#### Wiring

A single flexible coaxial cable (50 Ω, CW2040-3650SR, Cooner Wire) is used for power, communication, and image data transmission relying on a passive power-over-coax filter and a seriailzer/deserializer pair (TI DS90UB913A/DS90UB914A) for combining and separating DC power, low-speed bi-directional communication, and high-speed uni-directional imaging data. For coaxial cable lengths longer than 2.5 meters, an external 6V power supply should be connected to the UCLA Miniscope DAQ, replacing the power supplied by the USB connection to the MiniLFOV. With a total cable diameter down to 0.3 mm (Molex 100065-0023) and compatibility with active and passive, low torque commutators, this design minimizes the impact of cabling on animal behavior. The DAQ hardware and software are based on the UCLA Miniscope project’s previous work (http://miniscope.org/index.php/Main_Page), with updated firmware (https://github.com/Aharoni-Lab/Miniscope-DAQ-Cypress-firmware) and software (https://github.com/Aharoni-Lab/Miniscope-DAQ-QT-Software) to enable video streaming and controlling of the MiniLFOV. This updated software enables excitation LED brightness adjustment, focus adjustment by EWL, real-time ΔF/F and fluorescent trace visualization, frame rate selection, gain adjustment, and supports real-time pose estimation using DeepLabCut-Live^50^ through embedded Python. A list of the hardware and software used is given in **Table 3**.

### Animals

All experimental protocols were approved by the Chancellor’s Animal Research Committee of the University of California, Los Angeles, in accordance with the US National Institutes of Health (NIH) guidelines.

### Surgical implantation

#### Rat hippocampal imaging

3-month-old Long-Evans rats (*Charles River*) underwent two survival surgeries prior to behavior training to record fluorescent Ca^2+^ activity from hippocampal CA1 cells. During the first surgery, rats were anesthetized with 5% isoflurane at 2.5 L/min of oxygen, then maintained at 2-2.5% isoflurane while a craniotomy was made above the dorsal hippocampus. Next, 1.2 uL of AAV9-Syn-GCamp7s (AddGene) was injected just below the pyramidal layer (−3.6 AP, 2.5 ML, 2.6 DV) via a 10 uL Nanofil syringe (World Precision Instruments) mounted in a Quintessential Stereotaxic Injector (Stoelting) controlled by a Motorized Lab Standard Stereotax (Harvard Apparatus). Left or right hemisphere was balanced across all animals. One week later, the rat was again induced under anesthesia and 4 skull screws were implanted to provide stable mounting for the GRIN lens implant and MiniLFOV baseplate. The craniotomy was reopened to a diameter of 1.8 mm, and cortical tissue and corpus callosal fibers above the hippocampus were aspirated away using blunted 27 and 30 gauge needles. Following this aspiration, and assuring no bleeding persisted in the craniotomy, a 1.8 mm-diameter GRIN lens (#64-519, Edmund Optics) was implanted over the hippocampus and cemented in place with methacrylate bone cement (Simplex-P, Stryker Orthopaedics). The dorsal surface of the skull and the bone screws were cemented with the GRIN lens to ensure stability of the implant, while the surface of the lens was left exposed. Two to three weeks later, rats were again placed under anesthesia to cement a 3D printed baseplate above the lens. First a second GRIN lens was optically glued (Norland Optical Adhesive 68, Edmund Optics) to the surface of the implanted lens and cured with UV light. The pitch of each GRIN lens was approximately 0.25, so combining 2 in series provided roughly a 0.5 pitch. This half pitch provides translation of the image at the bottom surface of the lenses to the top while maintaining the focal point below the lens, effectively becoming a relay GRIN lens. This relay implant enables access to tissue deep below the skull surface. It has been simulated that the aberrations of the 1.8-mm GRIN near the edges generate a rotational stretching of the neural footprints which can be alleviated with high-quality GRIN lenses (provide roughly a 90% clear aperture) or commercially available lower NA GRIN lenses with near full FOV image achievable. Cannula (2.5 mm to 3 mm in diameter) was also reported to be used for imaging hippocampal CA1 instead of GRIN lens^51–54^. The MiniLFOV was placed securely in the baseplate and then mounted to the stereotax to visualize the Ca^2+^ fluorescence and tissue. The baseplate was then cemented in place above the relay lenses at the proper focal plane, the MiniLFOV was removed from the baseplate, and cement allowed to cure. Once rats had been baseplated, they were placed on food restriction to reach a goal weight of 85% ad lib weight and then began behavioral training. Briefly, the MiniLFOV was mounted into a baseplate with two set screws (4-40 × 1/4”), then positioned above the implanted lenses while the baseplate was cemented in place with bone cement (**Fig. 3b**). The baseplate body was 3D printed (20 mm × 20 mm outer dimensions) with 2 set screws on the side (**Fig. 1a**, **Fig. s6–s7**) for rigidly mounting a protective cap when not recording (protecting the GRIN lens and surgical region) and MiniLFOV body during experiments. *Rat histology*. At the end of the experiment, rats were anesthetized with isoflurane, intraperitoneally injected with 1 mL of pentobarbital, then transcardially perfused with 100 mL of 0.01M PBS followed by 200 mL of 4% paraformaldehyde in 0.01M PBS to fix the brain tissue. Brains were sectioned at 40 μm thickness on a cryostat (Leica), mounted on slides, then imaged on a confocal microscope (Zeiss) to confirm GFP expression and GRIN lens placement.

#### Cranial window imaging on head fixed mice

Adult C57BL/6N Tac (3-5 months old) male mice were singly housed on a 12 h light/dark cycle. Mice were bilaterally microinjected with 500 nL of AAV1.Syn.GCaMP6f.WPRE.SV40 virus (purchased from Addgene 100837) at 20 - 120 nL/min into the dorsal cortex using the stereotactic coordinates: −1.7 and −2.3 mm posterior to bregma, 0.5 mm lateral to midline and −0.8 mm ventral to skull surface. Mice underwent window implantation surgeries as previously described^55^. Briefly, a square region of skull ~4 mm in width was marked using stereotactic coordinates (center at bregma −2.2 mm AP). The skull was thinned using a dental drill and removed. After cleaning the surgical site with saline, a custom cut sterilized coverslip (square, 4 × 4 mm) was placed on the dural surface and fastened with adhesive and dental acrylics to expose a square window of approximately 3.5 mm spanning the midline. Three weeks later, a small 3D printed baseplate was cemented onto the animal’s head atop the previously placed dental cement. In the second mouse, an aluminum bar with two threaded holes was attached to stabilize the mice during imaging sessions. Following baseplating or attachment of headbar, mice underwent handling (3 days) and habituation (3 days) to acclimate them to the treadmill and head-fixation.

### Rats Behavioral Training

#### Ca^2+^ imaging in freely behaving rats

After baseplating, rats were given 15-minute sessions every 48 hours to perform a linear alternation task on a rectangular linear maze where one arm (2.5 m) directly connects the reward locations, while the other 3 arms in total are indirect and, combined, twice as long (5 m). These arms form a 2.5 m × 1.25 m rectangle with the reward locations at the corners of one long side, elevated 1 meter off the ground. The paths are 10-cm wide spanned by small metal bars along the entirety of the short and long paths. The reward zones are coated wood, 14-cm wide, 20-cm long, at a 45-degree angle with the two paths. The entire maze has a short 1-cm wall to provide a safe graspable ledge in case the rat loses its footing. The outer perimeter of the short path has a slightly taller wall angled 45 degrees away from the path to protect the wiring connected to the short path bars. 20-mg chocolate sucrose pellets are delivered through a metal tube at the end of the reward zone via an automated hopper, controlled by a computer running Cheetah software (*Neuralynx*) feeding position information to custom MATLAB scripts which controlled pellet delivery as the rat ran from one reward zone to the other. Rewards are delivered at the opposite reward location from their previous reward as the rat passes through the center of either path. Both paths yield delivery of 2 pellets, and there is no time out period after delivery. The rat must enter the most recent reward delivery location before the opposite side will be rewarded, forcing an alternation behavior. The rats are allowed to choose either path, and since they are equally rewarded, they demonstrate a preference for the short path (2.5 m arm) over the long path (the other 3 arms) after only a few sessions. Rats are given 15-minute recording sessions every 48 hours to minimize potential photobleaching of the Ca^2+^ indicator. Once a rat demonstrated a consistent short path preference and sufficient running behavior (>2 short paths per minute and a 2:1 short:long ratio in the first 10 minutes for two consecutive sessions) we introduced a manipulation in the subsequent session. Effects of these manipulations are omitted in this article, and we only evaluate period of consistent behavioral performance in absence of induced learning.

#### Wire-free imaging on freely behaving rats

For testing the functionality of the wire-free configuration, the experiment was conducted by putting the rat into a circular open-field (80-cm diameter) to record the unconstrained exploration of the environment and the neural activity simultaneously. An additional red LED is attached onto the MiniLFOV body for the behavior camera to track its position. Animals were food-deprived up to 85% of their original weight after which recordings commenced. Sucrose grain pellets (20 mg) were thrown in the enclosed environment every 20 seconds at random locations within the open field, keeping the animal in continuous locomotion, thereby allowing close to complete sampling of the environment^56^.

### Data analysis

#### Behavior video analysis

For experiments involving freely behaving rats, videos of behaviors were captured in AVI format using an overhead camera (30 FPS, Logitech C270). The position of the rat is extracted by tracking the position of the red LED attached to the top of the MiniLFOV recorded by a behavioral camera. The pixel location of the LED is found by detecting the highest pixel value region within the gray-scale behavior recording frames, a correction of the optical aberration of the camera lens is applied, and then the pixel value location is converted to real-world coordinates. For the head-fixed experiment in mice, a MiniCAM (**Fig. s12a**) was used for recording movement speed of the mice. The movement of the mice was extracted from the recorded motion of a circular treadmill (22.9-cm diameter) marked with 1-cm wide alternating dark and white bars on the surface (**Fig. s12b**).

#### Head orientation analysis

The quaternion values (*q_w_, q_x_, q_y_*, *q_z_*), read from the on-board head orientation sensor (BNO055, Bosch) following each frame acquisition, were saved in CSV file format during recording. This data was converted to Euler angles (roll, pitch, yaw) offline for further analysis, with the matrix used below:

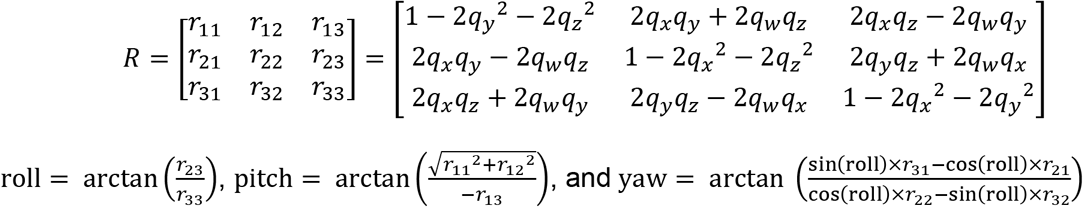

Further correction of the Euler angles depends on the axis orientation of the head-orientation sensor and the mounting orientation of the MiniLFOV during experiments.

#### MiniLFOV data preprocessing

The imaging data from the CMOS image sensor is saved in AVI file format using an uncompressed (GREY) video codec directly supported within the Miniscope DAQ software, and all the recorded videos for each animal in each session were concatenated into one single video file and cropped to the ROI pixel region using custom Python 3 scripts before further processing.

#### Wired experiment on rats in rectangular linear maze

Image stacks of Ca^2+^ dynamics (972 × 1296 pixels) were cropped to the pixel region containing the relay lens stack (720 × 720 pixels). We next used the CaImAn Python processing pipeline to perform non-rigid motion correction followed by image source extraction using constrained nonnegative matrix factorization for endoscopic recordings (CNMF-E)^27,28^ to identify and extract the spatial contours and fluorescent Ca^2+^ activity of individual neurons. Fast non-negative deconvolution OASIS (https://github.com/zhoupc/OASIS_matlab) was used to deconvolve the slow time course inherent in the GCaMP fluorophore^28^ to estimate underlying neural activity. The resulting deconvolved Ca^2+^ activity can be interpreted as the sum of neural activity within each frame scaled by an unknown constant, and we refer to this measure as the ‘temporal neural activity’ of a cell. Because the temporal neural activity of each cell is scaled by an unknown number that varies across cells, we normalized each cell’s temporal neural activity. Spatial neural activity rates were calculated path by path (Path 1 to 8) using 2-cm-wide spatial bins and a speed threshold of greater than 20 cm/s. Temporal neural activity and occupancy of the animal were spatially binned and then smoothed using a Gaussian kernel with σ=5 cm. Then the binned neural activity was divided by the binned occupancy to calculate the spatial neural activity rate of each cell.

The information content of the spatial neural activity rate map (in bits)^17^ of a single neuron was defined as:

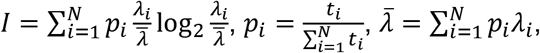

where *t_i_* represents the occupancy time spent in the *i*-th bin of total *N* bins and *λ_i_* represents the neural activity rate in the *i*-th bin. 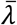 is the mean neural activity rate. The significance of the spatial information was calculated using two circular shuffling procedures. For each shuffling procedure, the random permutations were generated by frame-shifting the speed-thresholded sequence of temporal neural activity for each trail (total 32 trials) by a random interval between 0 and the number of frames for each trial respectively with frame number out of range being circularly wrapped to the beginning^57^. Then random permutations were generated by frame-shifting the entire sequence of temporal neural activity along the rat’s 8 paths by a random interval between 0 and the number of frames, with frame number out of range being circularly wrapped to the beginning. This procedure was repeated 500 times with random shifts to determine a significance measure for the spatial activity rate map of each neuron. The significance measure is the percentile of the true information content value within the distribution of the information content values from the 500 randomly shuffled sessions. Cells information content significantly above chance (*P*≥0.95) based on the circularly shuffled distribution are labelled place cells.

#### Head-fixed experiment imaging dorsal cortex in mice

As described earlier, Ca^2+^ recordings were processed as to yield the spatial contours of neurons and their deconvolved Ca^2+^ activity. Distribution of cell numbers is calculated inside 500 randomly chosen 1-mm diameter circular regions (FOV of Miniscope V4) in the entire FOV of the MiniLFOV.

#### Wire-free imaging on freely behaving rats

Wire-free MiniLFOV data was extracted from microSD cards and saved as uncompressed 8 bit AVI video files using custom Python code for processing and analysis. Ca^2+^ recordings were processed as described earlier to yield the spatial contours of neurons and their deconvolved Ca^2+^ activity. Spatial neural activity rates were calculated using 2 cm × 2 cm spatial bins and a speed threshold of greater than 5 cm/s. Temporal neural activity and occupancy of the animal were spatially binned and then smoothed using a Gaussian kernel with σ=3 cm. A minimum occupancy threshold was set to be 100 ms where spatial bins that did not meet this threshold were excluded from all subsequent analysis. The binned neural activity was divided by the binned occupancy to calculate the spatial neural activity rate of each cell. Information content of the spatial neural activity rate map was calculated with the same equation as described in the wired experiment on rats in the rectangular track and the significance of the spatial information content was calculated using a circular shuffling procedure. Random permutations were generated by time-shifting the entire sequence of positions along the rat’s path by a random interval between 10 s and the total recording minus 10 s, with the time out of the range being wrapped to the beginning^57^. The 10-seconds at the beginning and the end of the recoding are occluded to remove the period of putting the rats into and out of the open-field. This procedure was repeated 500 times to determine a significance measure for the spatial activity rate map of each neuron. The significance measure is the percentile of the true information content value within the distribution of the information content values from the 500 randomly shuffled sessions. Cells with information content significantly above chance (*P*≥0.95) based on the circularly shuffled distribution are labelled place cells. For cells defined as place cells, their place fields are defined as the region that 1) contains 5 adjacent bins with binned neural activity rate above 95% of that of all randomly shuffled binned neural activity rates and 2) extends to all connected bins with a binned neural activity rate of at least 5% of the place field’s peak binned neural activity rat. This approach allows for a robust detection and spatial region definition for place cells with single as well as multiple place fields. The place field spatial region is shown as red contours in **Figure 4h**.

### Matching cells across sessions

Contours computed from CaImAn were normalized by their own maximum value then thresholded at 0.5 to generate compact contours for all cells. Artifact contours detected on the edge of the GRIN lens were manually removed then each session was manually aligned using the contours and vasculature simultaneously using custom MATLAB scripts. In addition to linear translations, it was sometimes necessary to radially scale this data as well to account for slight differences induced by camera placement and different focal positions from the electrowetting lens. For all rats, we aligned and matched 3 consecutive recording sessions (48 hours apart) where running performance was stable. Contour alignment was performed using CellReg to register cells across all sessions of interest based on their centroid distances and spatial contour correlations^58^. Distributions of centroid distances and spatial correlation for all cell pairs within 24 μm were computed, yielding a two-dimensional distribution that can be modeled and given a probability threshold to match cell pairs. Cell pairs with a probability >0.5 for both the centroid and correlation distributions were matched (low centroid distance and high spatial correlation), which is expected when recording the same cell.

## Data availability

The experimental data that support the findings of this study are available from H. Tad Blair, Alcino J. Silva, and Daniel Aharoni upon reasonable request.

## Code availability

Data analysis scripts are available on reasonable request from H. Tad Blair, and Daniel Aharoni.

## ACKNOWLEGMENTS

Changliang Guo was supported by the National Institute of Mental Health (1UF1NS107668).

Garrett Blair was supported by NeuroNex grant, National Science Foundation (award # 1707408).

Megha Sehgal was supported by R01 MH113071, Adelson Adelson Medical Research Foundation and DBI-1707408.

Federico N.Sangiuliano Jimka was supported by NeuroNex grant, National Science Foundation (award # 1707408).

Arash Bellafard was supported by NIH Grant 1R01NS116589-01.

## AUTHOR CONTRIBUTIONS

Daniel Aharoni, Changliang Guo, H. Tad Blair, Michele A. Basso, Peyman Golshani, and Alcino J. Silva conceived the study design.

Daniel Aharoni, and Changliang Guo developed the MiniLFOV.

Federico N.Sangiuliano Jimka contributed to dual-LED circuit design.

Garrett Blair performed the surgeries in rats.

Garrett Blair, and Changliang Guo conducted the rat experiments.

Megha Seghal performed the cranial window surgeries in mice.

Megha Sehgal, Changliang Guo, and Garrett Blair conducted the head-fixed experiments on mice.

Arash Bellafard contributed to testing MiniLFOV on head-fixed mice.

Changliang Guo and Garrett Blair performed data analysis.

Changliang Guo, Garrett Blair, Megha Sehgal, and Daniel Aharoni prepared the manuscript and figures.

## Supplemental Information

### Miniscope-LFOV

A large field of view, single cell resolution, miniature microscope for wired and wire-free imaging of neural dynamics in freely behaving animals.

**Video 1.**
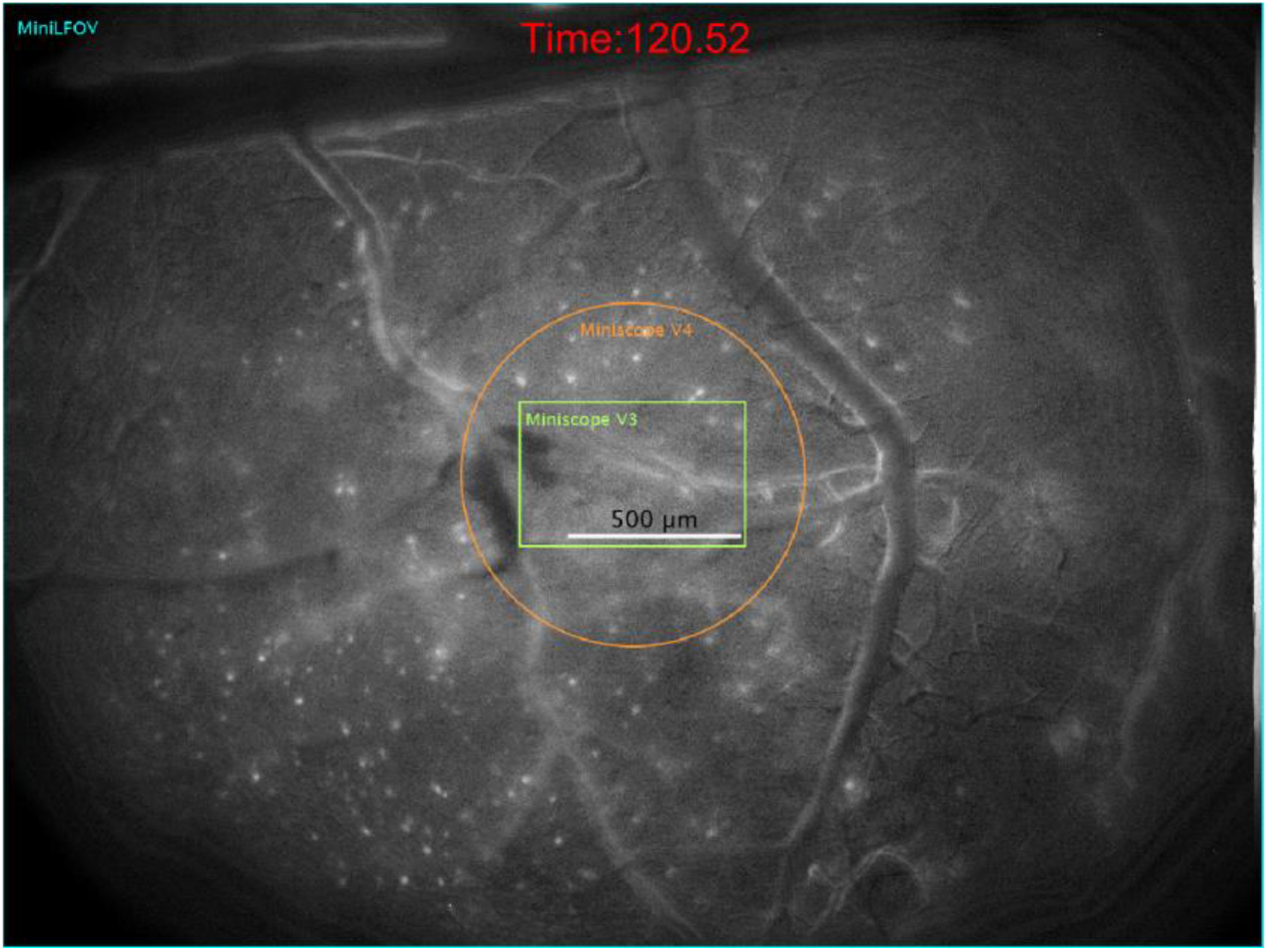
FOV of Miniscope V3, Miniscope V4, and MiniLFOV. MiniLFOV has 30-fold increase in FOV than the previous generation UCLA Miniscope V3 and 12-fold increase in FOV than the current generation UCLA Miniscope V4. The background was calculated as the minimum projection of the raw video (after motion correction) and removed to show clear neuronal activities in this video.

**Video 2.**
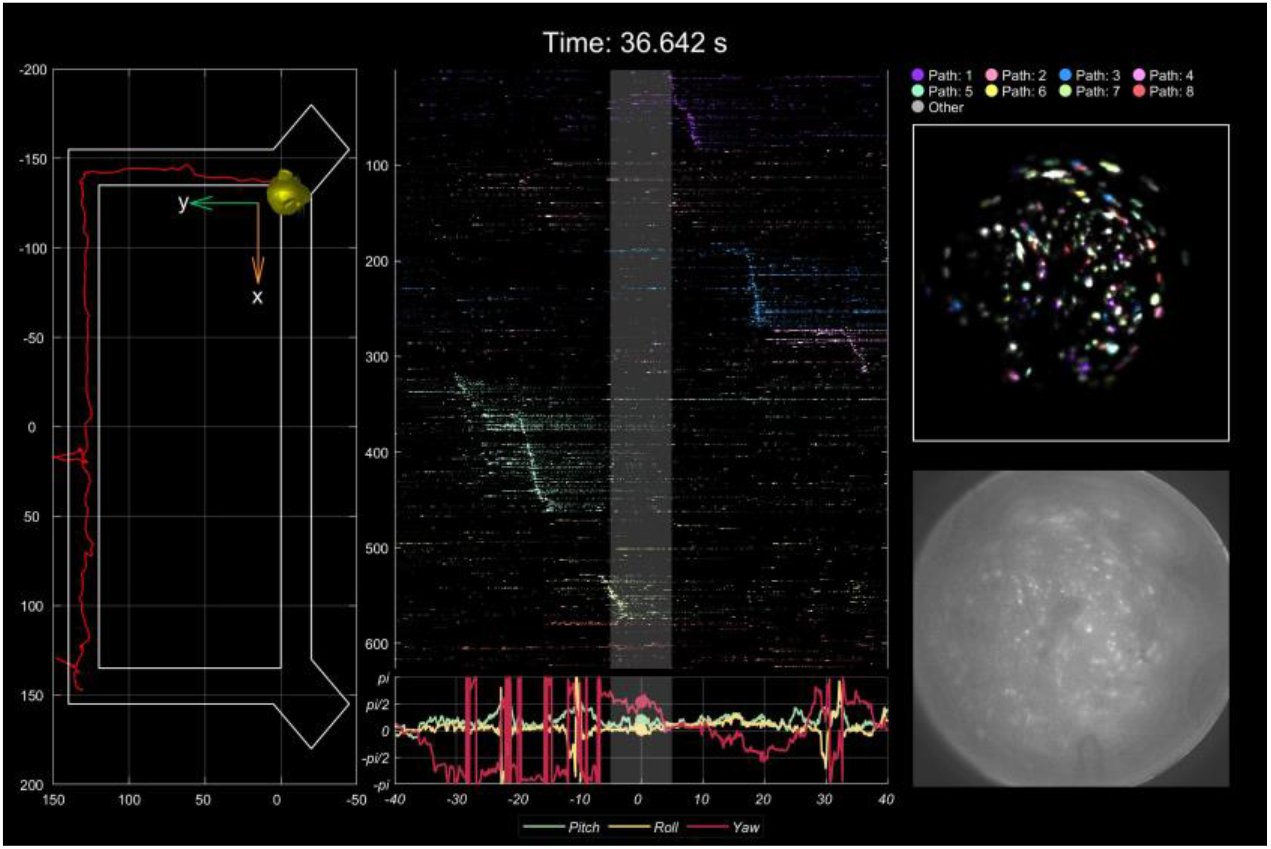
An example video showing the position of the rat, head orientation, and place cell activity during behavior. To record place cell activity from the dorsal CA1 region in rats, we imaged Ca^2+^ dynamics in GCaMP7s-expressing neurons in dorsal CA1 while rats performed navigation of a rectangular track environment. We present one exemplary session of one rat to demonstrate the effectiveness of the system. Recordings took place on a rectangular track (2.5 m × 1.25 m), with two reward feeders on two corners of the track. Positions on the track and head orientation of the rat were extracted from a software synchronized behavioral camera and on-board head orientation sensor respectively. The raw video is 972 × 1296 pixels after within-sensor 2x pixel binning and then further cropped to the pixel region containing the relay GRIN lens (720 × 720 pixels). In this example recording, a total number of 1357 cells can be detected and extracted using CNMF-E analysis via CaImAn. As expected, a significant population of the cells (626; 46.2%) are spatially tuned along at least one direction on the 4 arms on the rectangular track. The behavior of the rat was serialized into 8 paths (4 arms with 2 running directions). Place cells recorded during the session were significantly modulated by both position and direction of travel and span across all regions of the track (**middle**). The raw video after motion correction shows as a reference (**bottom right**) for the video showing the activity of the extracted cells (**top right**, marked in 9 colors to represent the places cell for each path and other cells with spatial information below 95% of 500 randomly shifted deconvolved neural activity for each cell). These data demonstrate how the MiniLFOV can improve experimental efficacy by yielding more cells in larger research models, broadening the horizon of potential hypotheses for researchers.

**Video3.**
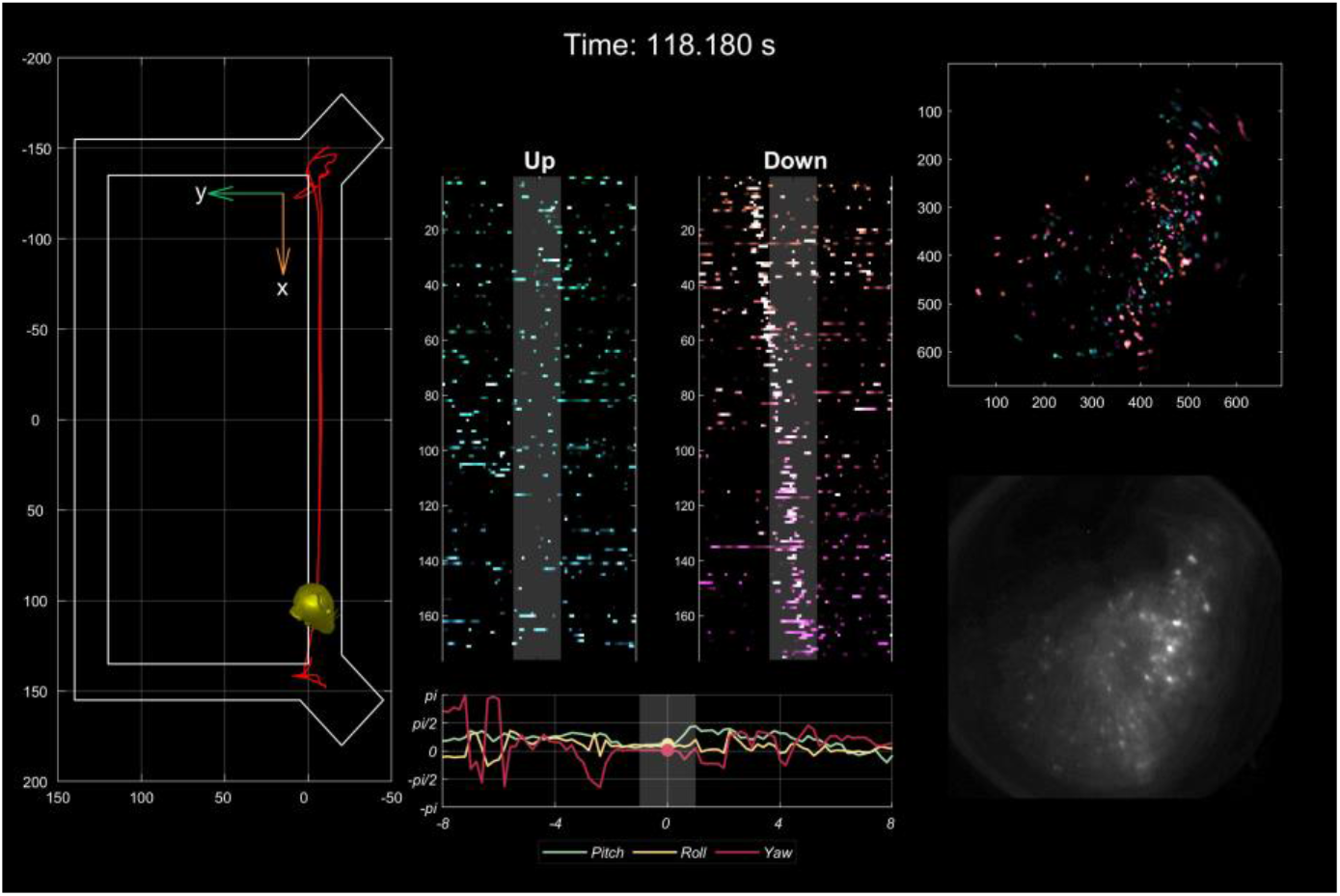
An example video showing the position of the rat, head orientation, and place cell activity during behavior. In this video, only two directions (Up and Down) on the long arm with two reward feeders on two corners are considered and plotted. Place cells related to running up and down are shown in the middle of the video.

**Figure s1.**
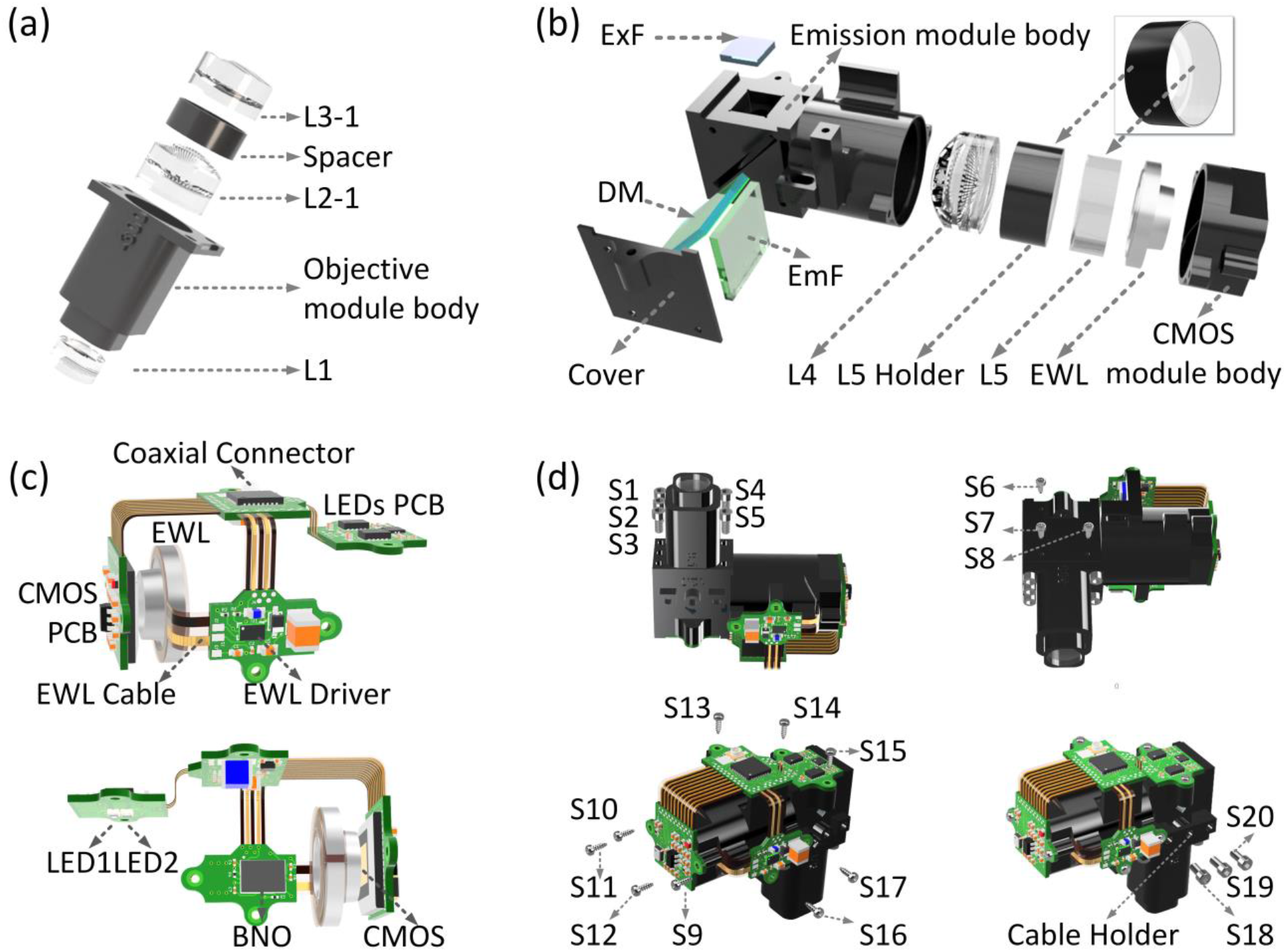
Mechanical and optical assembly of MiniLFOV. The MiniLFOV consists of three main modules, objective module (details in **a**), emission module (details in **b**), sensor module, and a custom Rigid-Flex printed circuit board (PCB) (details in **c**), and they are screwed together with two types of screws (details in **d**). Objective module (1.8-mm WD) contains three achromatic lenses (L1, L2-1, and L3-1) with a spacer (3.5-mm tall) placed between L2-1 and L3-1, with a 3D printed objective module body to hold the components. 3.5-mm-WD objective module contains three achromatic lenses (L1, L2-2, and L3-2) with a spacer (2-mm tall) placed between L1 and L2-2. Emission module contains filters (excitation filter “ExF”, dichroic mirror “DM”, and emission filter “EmF”), and optics lenses (achromatic L4 and plane-concave lens L5), with a 3D printed emission module body to hold the filters and lenses. Sensor module holds the EWL with a 3D printed CMOS module body. The circuit module consists of 4 sub-circuits, an excitation LED circuit, an electrowetting lens tuning and head orientation circuit, a CMOS image sensor circuit for collecting Ca^2+^ imaging data, and a power-over-coax and serializer circuit (**Fig. 1a, b**) for supporting coaxial cable power and data transmission. Two LEDs are housed on the LED circuit board driven by two LED drivers with I^2^C digital potentiometer for brightness adjustment. Electrowetting lens tuning and head orientation circuit consists of EWL driver, with EWL cable holding the EWL, to adjust the focus of the EWL and the absolute-orientation chip for collecting head orientation data. 5M CMOS sensor (MT9P031) is housed on the CMOS image sensor circuit for Ca^2+^ fluorescence image capturing. A serializer chip on the power-over-coax and serializer circuit serializes the imaging data and sends it over a coaxial connector to communicate with a custom MiniscopeDAQ system. The objective module, emission module, sensor module and the circuits module are assembled by 12 (S6-S17) M1 thread-forming screws (96817a704, McMaster-Carr), and 8 (S1-S5, S18-S20) 18-8 stainless steel socket head screws (0-80 Thread Size, 92196A052, McMaster-Carr).

**Figure s2.**
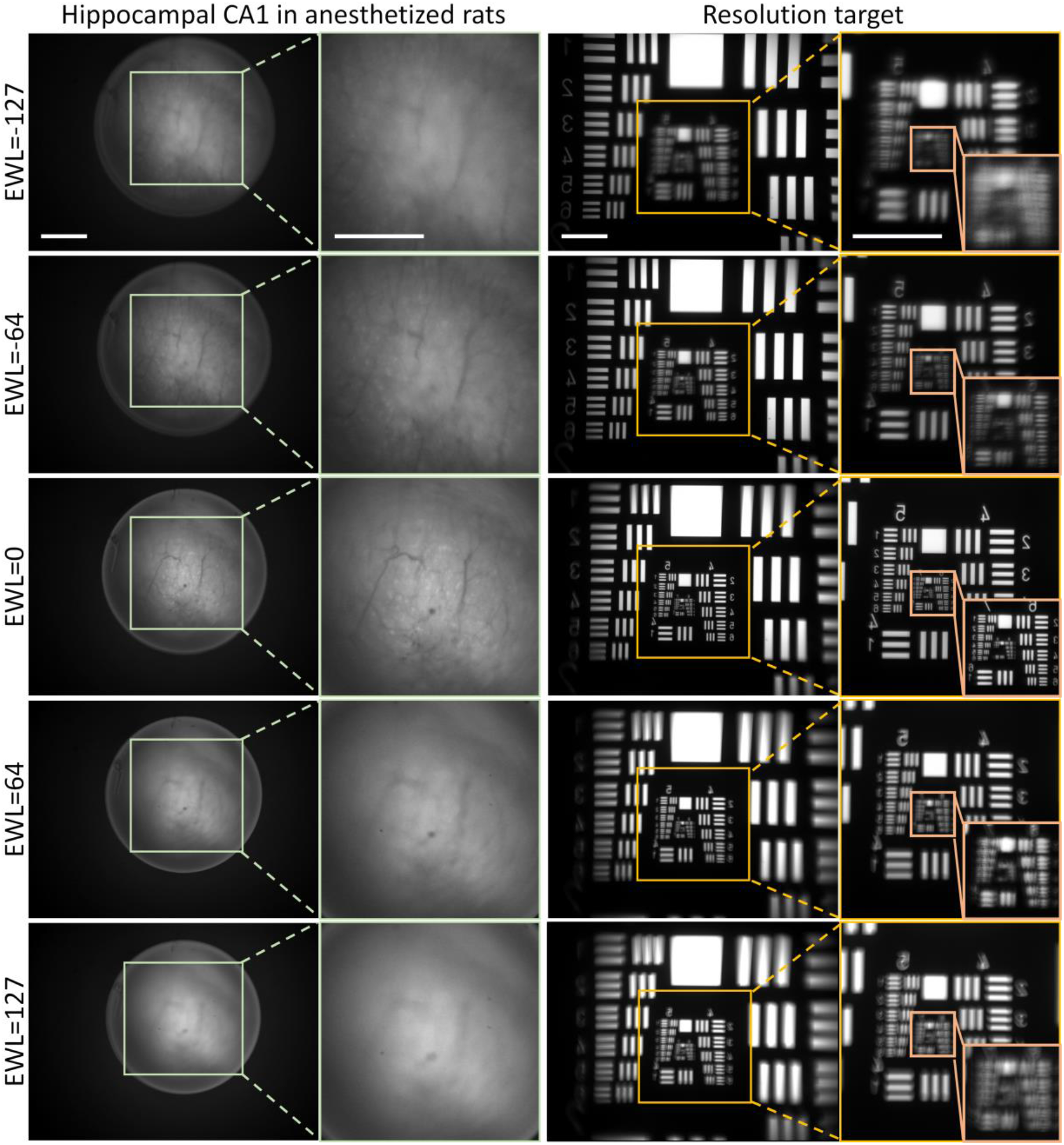
Five imaging focal planes attained by changing the focal position with the electrowetting lens (EWL) in an anesthetized rat and Resolution target to show how the resolution changes. The EWL value is set to be −127, −64, 0, 64, and 127 respectively. Negative EWL values relate to the deeper/ventral focal plane and positive EWL value corresponds to more superficial/dorsal. The cells and blood vessels are in focus when EWL is 0. The resolution of the MiniLFOV changes attributed to the change of the focal plane by adjusting the EWL, contributing to the resolution from 2.46 μm (Group 8 Element 5; 406 lps/mm) in focus (EWL=0) to 17.54 μm (Group 5 Element 6; 57 lps/mm) when EWL is set to be −127. Scale bar: 500 μm.

**Figure s3.**
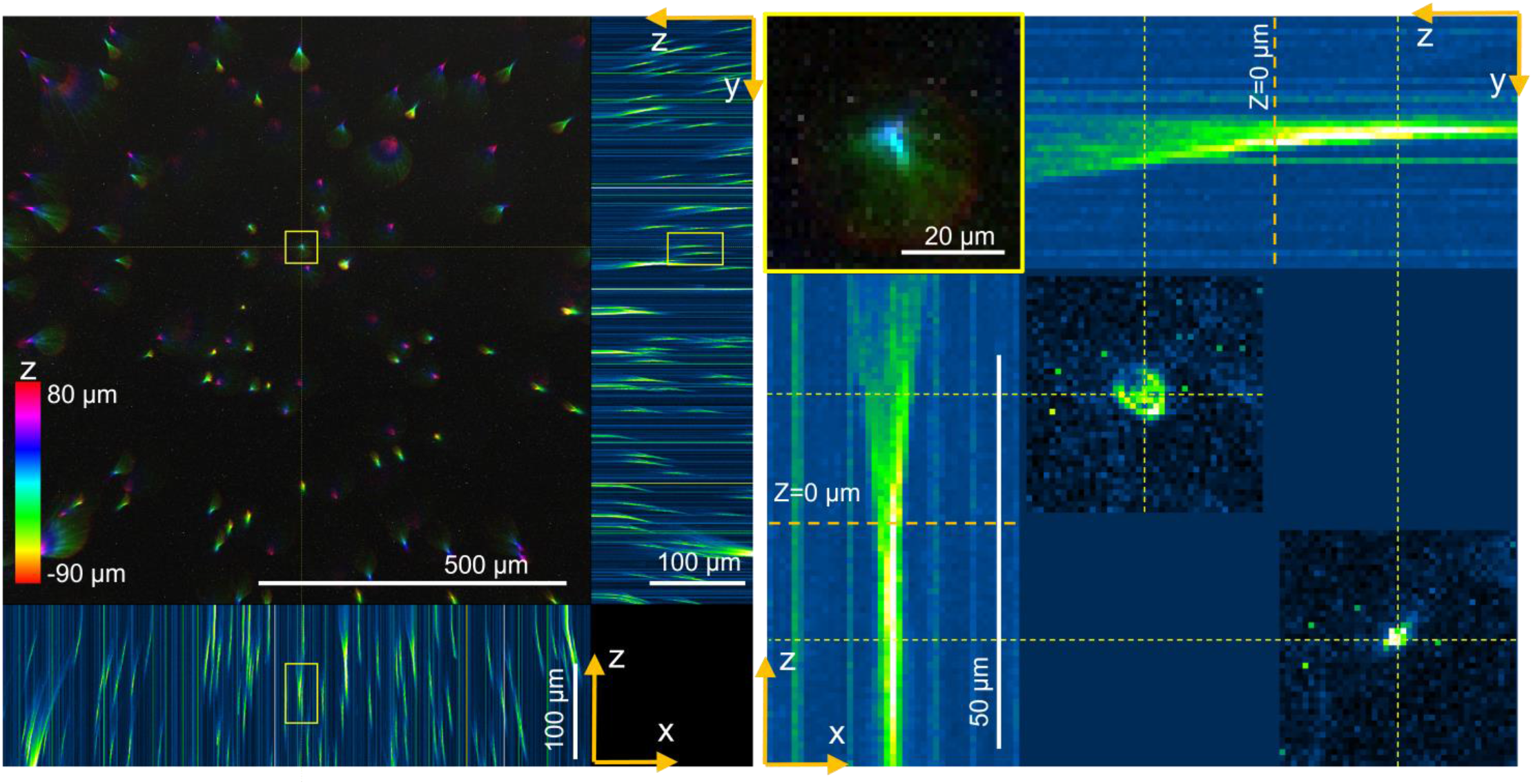
Imaging of 1-μm fluorescent beads in 2% agarose gel by changing EWL lens from −90 μm to 80 μm to show how the imaging changes. 0.5 μL of 1-μm fluorescent beads (10 μL diluted into 990 μL normal saline) are mixed with 500 μL 2% agarose gel. 40 μL of the agarose containing the fluorescent beads was applied to a coverglass (0.14 −0.17mm, 20×20 mm) and sealed onto a coverslide (1 mm thick) with UV glue to generate thick agarose gel with the beads randomly distributed inside, and the specimen was ready for use after 10 min. The sample slide is then attached to a slide holder (with the coverglass facing up). The MiniLFOV is attached to a manual 3D translation stage and placed above the sample slide. The lateral position and height of the MiniLFOV was adjusted to optimize image quality for each imaging session. Only the center 800 × 800 pixel (0.957 mm × 0.957 mm) region is shown in this figure. The left figure shows the color coded volumetric beads by adjusting EWL from −90 μm to 80 μm (with total 221 z-slices taken), with their projected x–z and y–z views shown on the bottom and right. Negative EWL values relate to the deeper/ventral focal plane and positive EWL value corresponds to more superficial/dorsal. The tilting of the beads along z axis is caused by the changed magnification of the imaging when adjusting EWL values. Distortions can be seen when the EWL value is near the adjustment limit. An example fluorescent bead located near focal plane (EWL=0) is shown in the right figure with x-z and y-z projections shown.

**Figure s4.**
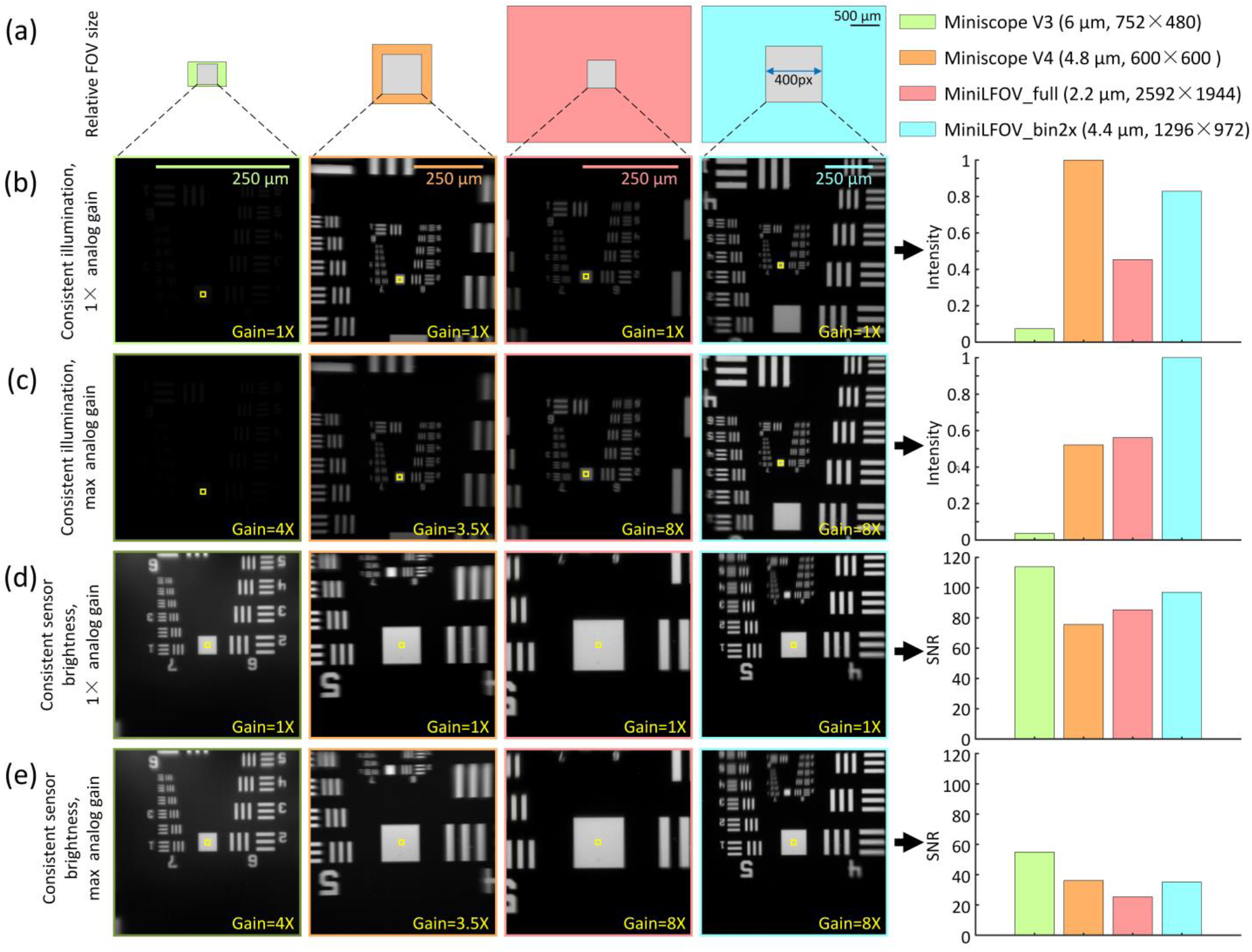
Sensitivity and signal noise ratio (SNR) comparison of Miniscope V3, Miniscope V4, MiniLFOV full resolution (MiniLFOV_full), and MiniLFOV after 2×2 pixel binning (MiniLFOV_bin2x) using Resolution Target. A light source was placed under a Negative USAF 1951 Hi-Resolution Target (#55-622, Edmund Optics) with a thin diffuser placed between them to diffuse the light. Then a Miniscope V3, Miniscope V4, and MiniLFOV are attached to a 3D translation stage and placed above the target. The lateral position and height of the Miniscopes were adjusted to optimize image quality for each imaging session. A 30-second video was recorded with each Miniscope in cases of consistent illumination (1×, and max analog gain), and consistent sensor brightness (1x, and max analog gain). The sensitivity comparison of the Miniscopes was done by illuminating the back side of the Negative USAF 1951 Hi-Resolution Target with the light source remaining at a constant intensity. Sensor sensitivity was calculated as the mean pixel value within the 10 x 10 pixel box shown in yellow. The SNR comparison was done by adjusting the light source intensity to achieve equivalent pixel brightness (8bit gray scale value = ~200), across all Miniscopes in the chosen area on the sensors. Sensor SNR was calculated as the mean pixel value divided by the standard deviation of all pixels within the 10 × 10 pixel box shown in yellow. A 400 × 400 pixel mean image from each 30-second recording is shown. **(a)** Relative FOV size of Miniscope V3 (light green box, 650 μm × 420 μm), Miniscope V4 (orange box, 1-mm diameter), MiniLFOV (salmon, and cyan, 3.1 mm × 2.3 mm), and chosen 400 × 400 pixel area (gray box) in each case. Legend shows pixel size and full sensor resolution. **(b)** Sensitivity of Miniscopes with unity analog gain set for each. MiniLFOV is less sensitive than Miniscope V4 (83%). **(c)** Sensitivity of Miniscopes with maximum analog gain set for each. Higher sensitivity is achieved with MiniLFOV than Miniscope in this case (192%). **(d)** SNR of Miniscopes with unity analog gain set for each. **(e)** SNR of Miniscopes with maximum analog gain set for each.

**Figure s5.**
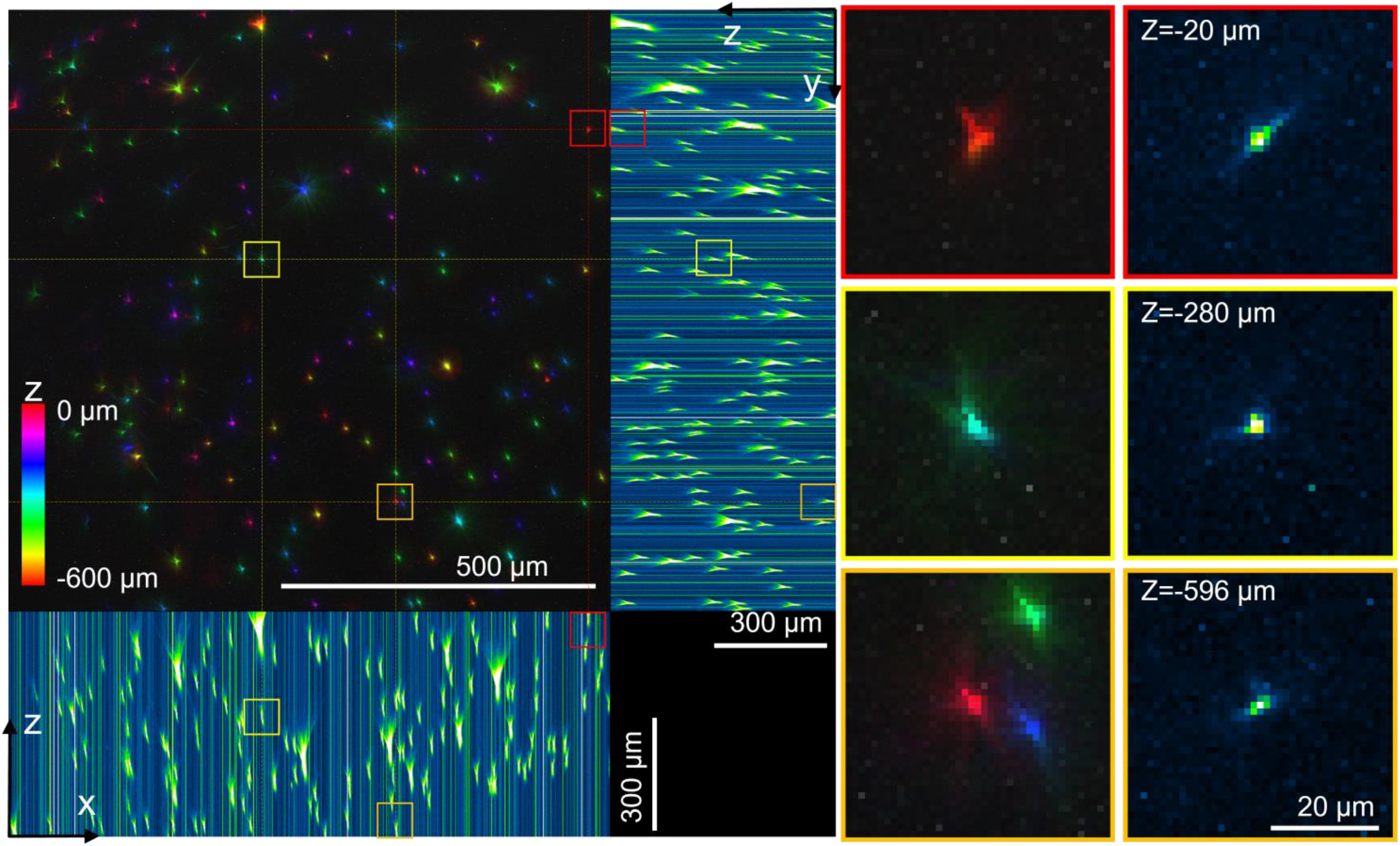
Imaging of 1-μm fluorescent beads up to 600 μm deep into thick 2% agarose gel sample slide. 40 μL of the agarose containing the fluorescent beads was applied to a coverglass (0.14 −0.17mm, 20×20 mm) and sealed onto a coverslide (1 mm thick) with UV glue to generate thick agarose gel with the beads randomly distributed inside, and the specimen was ready for use after 10 min. The sample slide is then attached to a slide holder (with the coverglass facing up) mounted on a motorized z-axis linear translation stage, to adjust the z-axis position of the sample with up to 100 nm adjustment accuracy. Then the MiniLFOV is attached to a manual 3D translation stage and placed above the sample slide. The lateral position and height of the MiniLFOV was adjusted to optimize image quality for each imaging session. The left figure shows the imaged color-coded beads distributed in 3D space in agarose gel by adjusting the z-axis translation stage closer to the MiniLFOV from a start imaging depth 0 μm (the depth close to the MiniLFOV) to −600 μm deep into the agarose gel, to make the MiniLFOV focusing deeper into the agarose gel step by step, with total 301 images taken for the stack. Their projected x–z and y–z views are shown on the bottom and right. Only the center 800 × 800 pixel (0.957 mm × 0.957 mm) region is shown in this figure. 3 example beads located around −20 μm (red box), −280 μm (yellow box), and −596 μm (orange box) are shown in the right figures to validate the MiniLFOV can resolve the fluorescent beads at least down to 600 μm (see the bead inside the orange box) into the agarose gel. Bright straight lines in the x–z and y–z projection figures are noises from low excitation light intensity.

**Figure s6:**
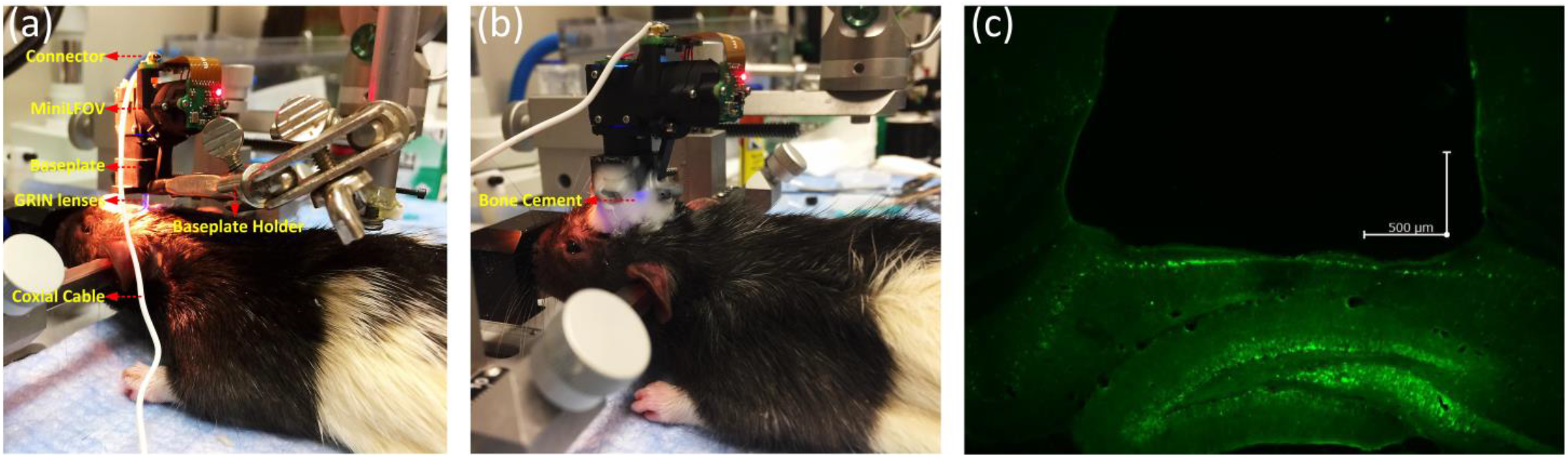
MiniLFOV baseplating on the rat. (**a**) A second ~0.25-pitch GRIN lens is affixed atop of the first ~0.25-pitch GRIN lens (implanted during earlier surgery), to construct a relay optics setup (~0.5 pitch). Then the MiniLFOV baseplate is held by a stereotaxic clamp. The MiniLFOV, attached to the baseplate with two screws, is positioned above the second GRIN lens on an anesthetized ear-barred rat. The distance from the MiniLFOV to GRIN lens is adjusted to focus on the hippocampal CA1 region exhibiting the most active and in-focus neurons. (**b**) The MiniLFOV baseplate is then affixed to the animal via bone cement mixture. After baseplating, a protective cap is used to cover and protect the GRIN lens from dust and scratches. (**c**) Histological image of a brain slice using a confocal microscope (Zeiss) sectioned at 40-μm thickness on a cryostat (Leica) to confirm GFP and GRIN lens placement. Picture shows GFP expression in the dorsal hippocampus of the rat, approximately 2.6 mm below the skull surface; scale bar: 500 μm.

**Figure s7.**
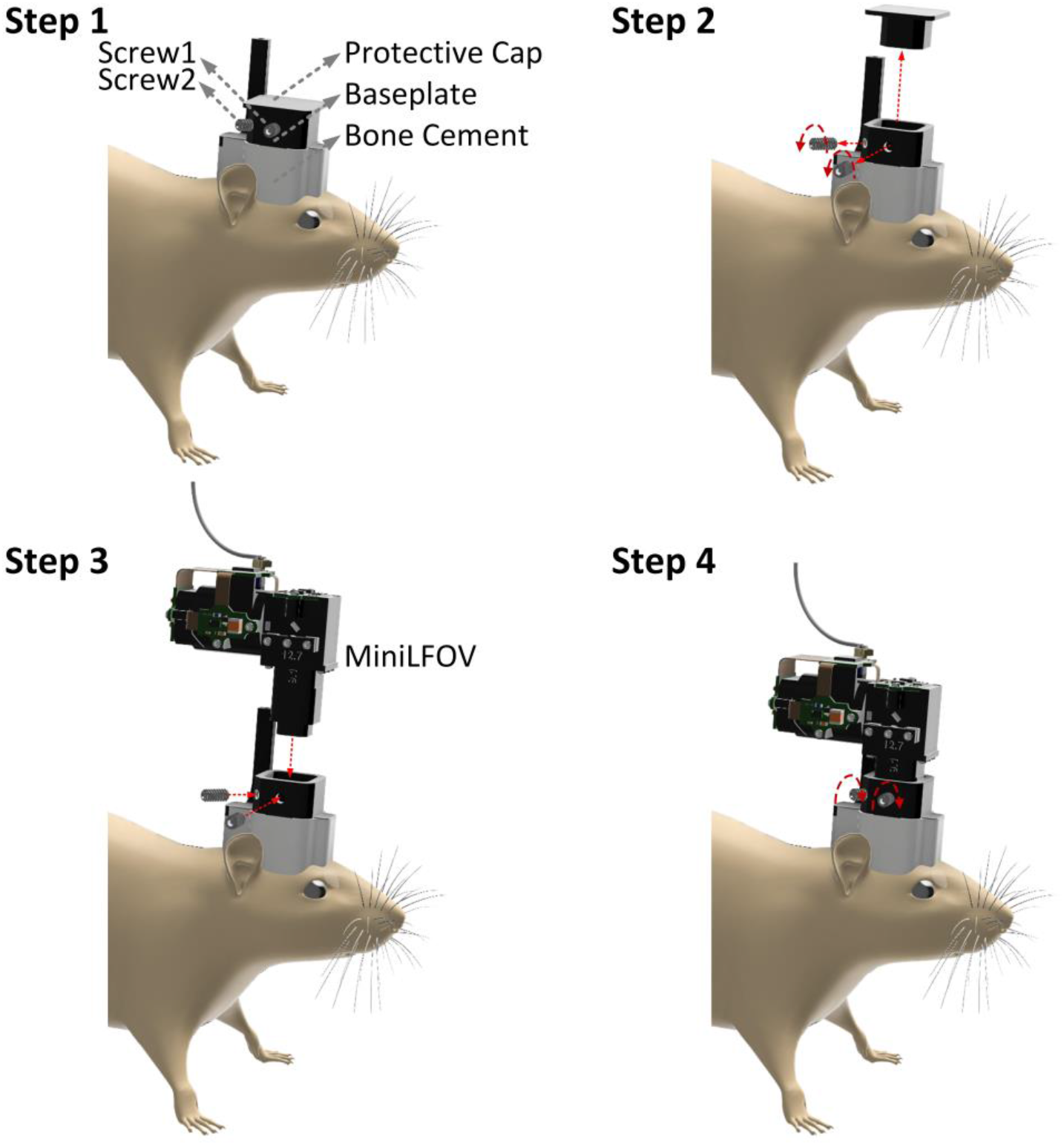
Mounting the MiniLFOV on the rat’s head. (**Step 1**) While the animal is in its home cage, a protective cap is used in the baseplate and secured with two set screws (4-40 × 1/4”) to protect the GRIN lens from dust and scratches. At the start of the experiment, the protective cap is removed (**Step 2**) and the MiniLFOV is inserted into the baseplate and secured with two set screws (**Step 3-4**). At the end of the imaging session, the MiniLFOV is again replaced by the protective cap.

**Figure s8.**
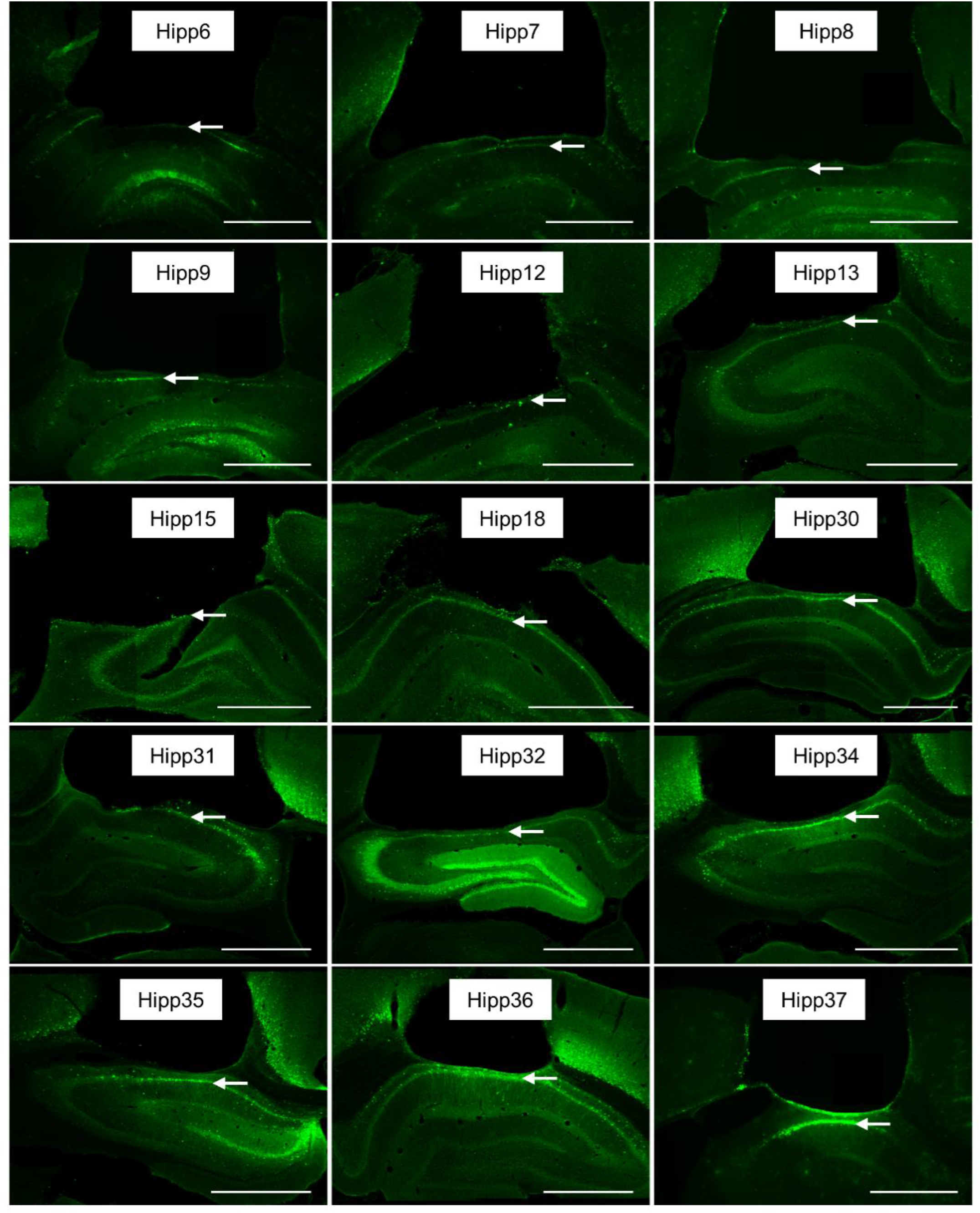
Example histology from 15 rats, showing GCaMP expression with the pyramidal layer for CA1 beneath the implanted lens location (arrow). Scale bar: 1 mm.

**Figure s9.**
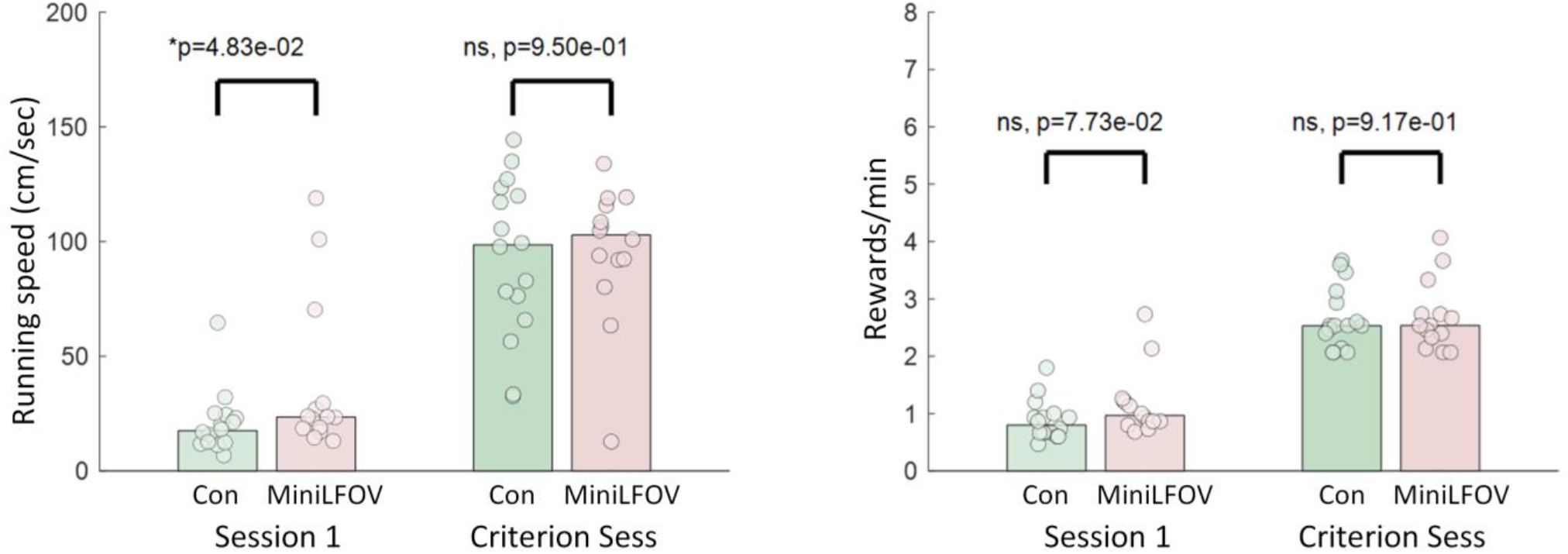
The test on the behaviors of the rats with and without MiniLFOV mounted. Both the first session and the session where they reach criterion for each group are shown in this figure. It can be seen that although the rats with MiniLFOV mounted start out running a bit faster than controls, they are never impaired compared to control in either measure. So the MiniLFOV doesn’t limit their movement ability.

**Figure s10.**
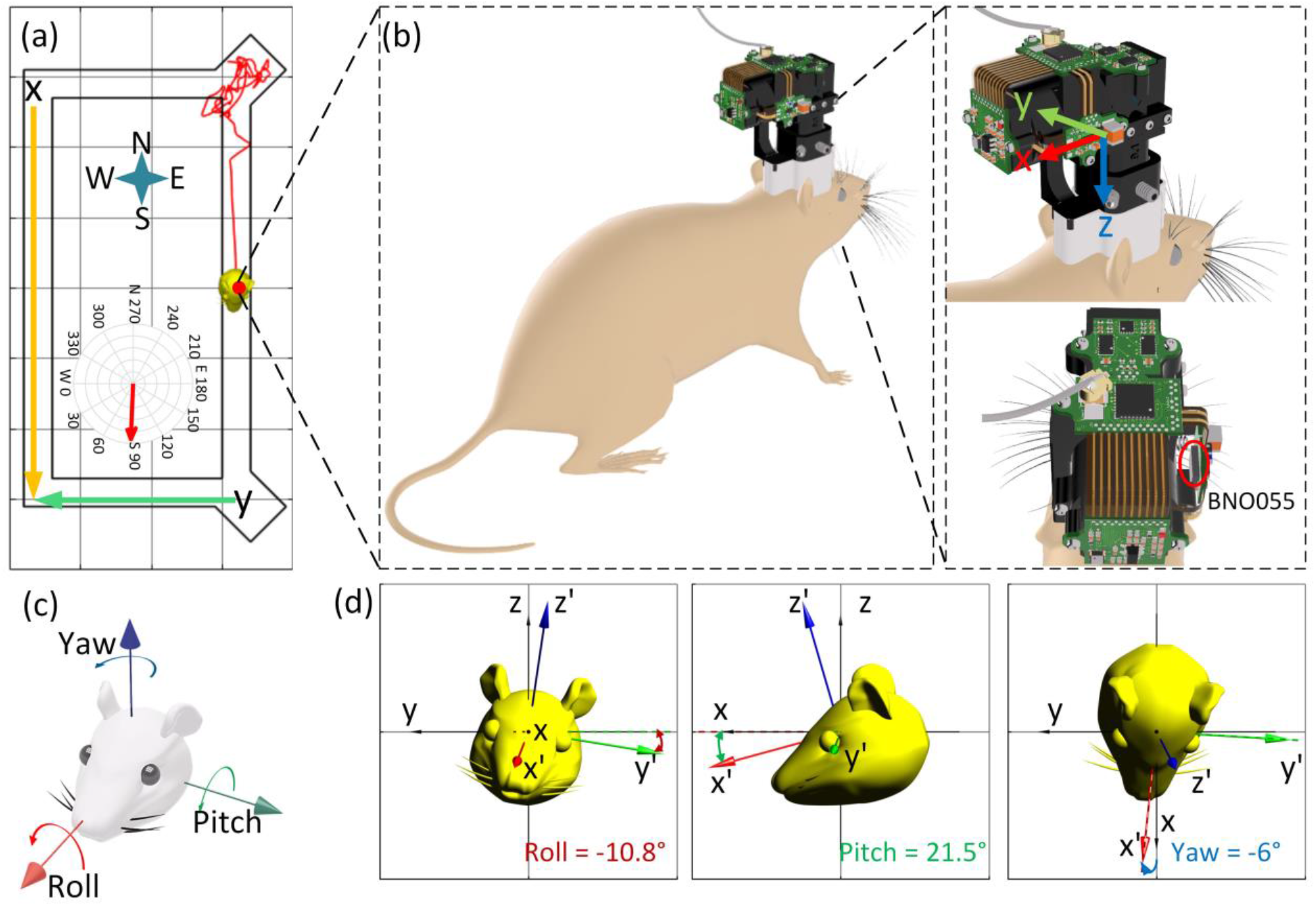
Illustration of the head orientation data (roll, pitch, yaw) from head-orientation sensor (BNO055). **(a)** A rat with a MiniLFOV mounted on its head running on a rectangular track. **(b)** 3D image showing the coordinate of BNO055 configured and its orientation. The axis orientation of the BNO055 can be re-configured to the new reference axis before experiment. **(c)** Illustration of roll, pitch, and yaw in terms of direction and rotation. **(d)** Zoomed-in figures shows the example roll (−10.8°), pitch (21.5°), and yaw (−6°) extracted from the recorded quaternion value (*q_w_*, *q_x_*, *q_y_*, *q_z_*), when the rat is passing x=0, running from north to the south on the track.

**Figure s11.**
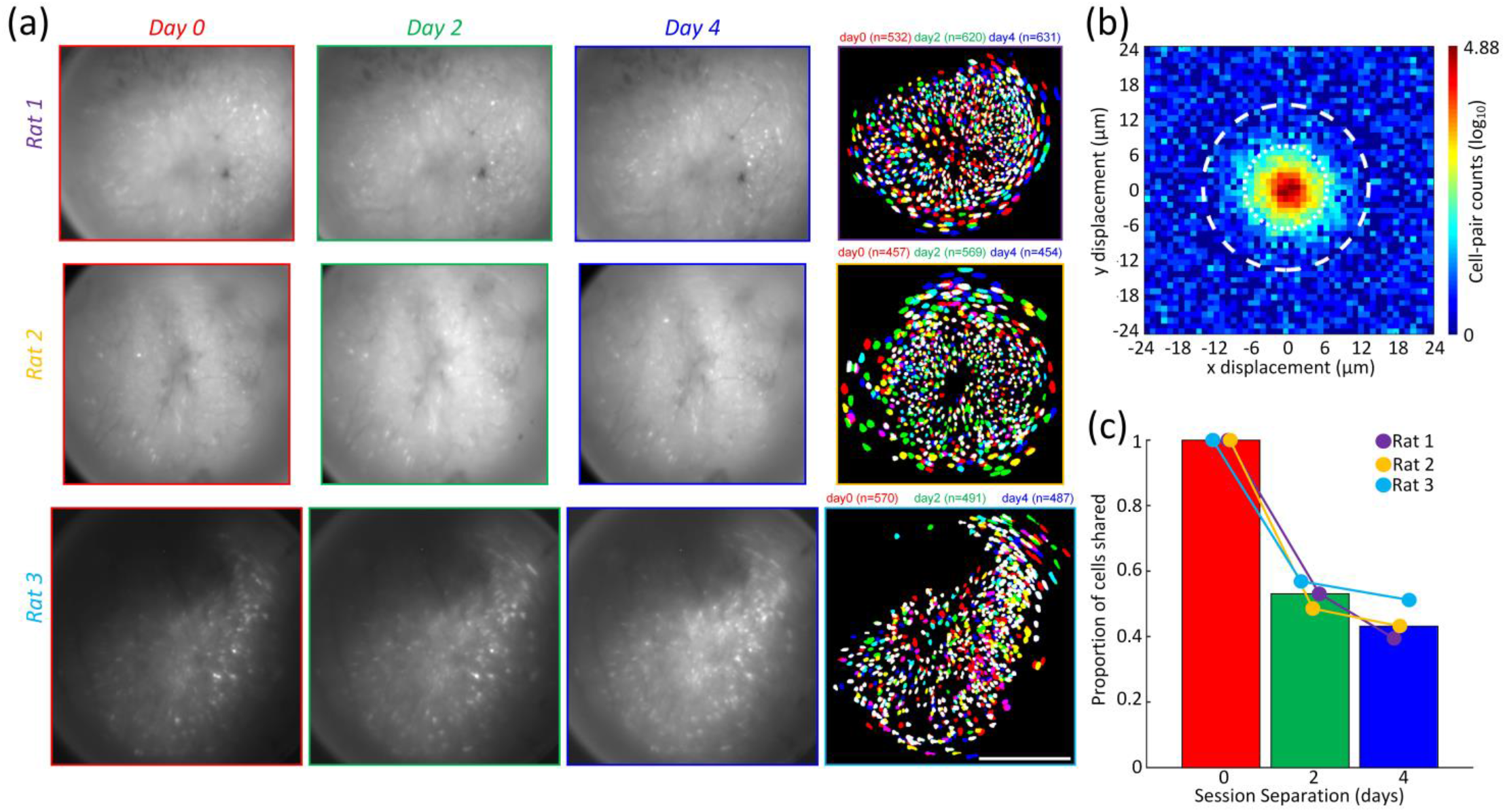
Cells can be reliably recorded across sessions. Three imaging sessions were performed in the same task with 48-hours separating each recording to demonstrate that the MiniLFOV is capable of consistently recording activity from the same population of cells across multiple sessions. **(a)** Mean frames (Column 1-3) and extracted contours of cells (Column 4) across all three recording days from Rat 1 (N_0_=532, N_2_=620, N_4_=631), Rat 2 (N_0_=457, N_2_=569, N_4_=454), and Rat 3 (N_0_=570, N_2_=491, N_4_=487). The contours of cells are colored by the recording day (day0=red, day2=green, day4=blue). The algorithm to match cells across all sessions is in **Methods**. Scale bar: 500 μm. **(b)** Distribution of centroid displacements for all cell pairs within a 24-μm radius, showing a large population of cell within the same location after across-day alignment. Distributions of centroid distances and spatial correlation for all cell pairs within 24 μm were computed, yielding a two-dimensional distribution that can be modeled and given a probability threshold to match cell pairs. Cell pairs with a probability >0.5 for both the centroid and correlation distributions were matched (low centroid distance and high spatial correlation), which is expected when recording the same cell. **(c)** Population overlap (the number of cells shared between sessions) the average population correlation vector between recording sessions decreased as a function of time from Rat 1 (day0=100%, day2=53.1%, day4=39.5%), Rat 2 (day0=100%, day2=48.6%, day4=43.2%), and Rat 3 (day0=100%, day2=56.9%, day4=51.2%).

**Figure s12.**
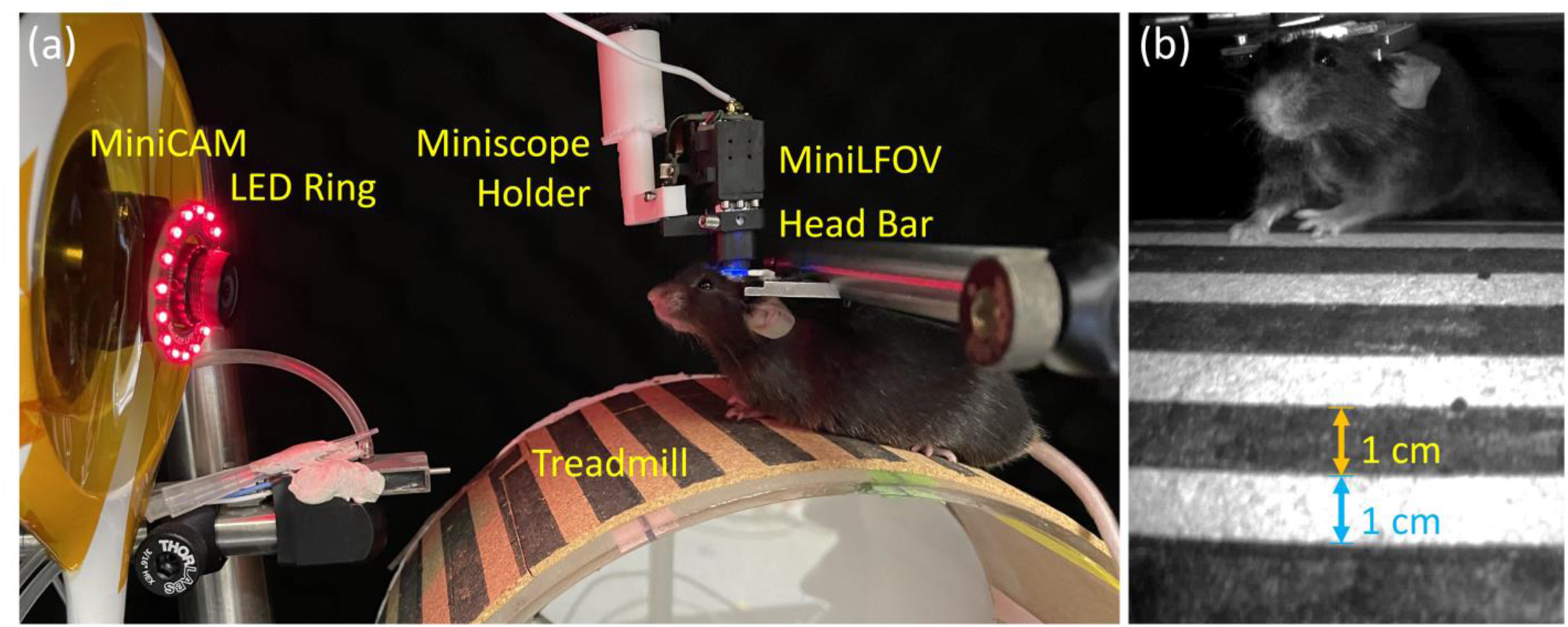
Experimental setup for Ca^2+^ imaging of dorsal cortex through a cranial window in head-fixed mice. **(a)** Experimental setup. The mice were head-fixed atop a circular treadmill (22.9-cm diameter and the white and dark bars with 1-cm wide). The MiniLFOV was attached to an articulated ball-head holder mounted on a 3D translation stage built of three linear translation stages (XR25C/M, Thorlabs) so that the lateral position and height of the MiniLFOV relative to the cranial window can be adjusted. A MiniCAM is used to capture the movement speed of the mice and the red LED ring is used for illumination in dark environment. **(b)** Single frame taken by the MiniCAM.

**Figure s13.**
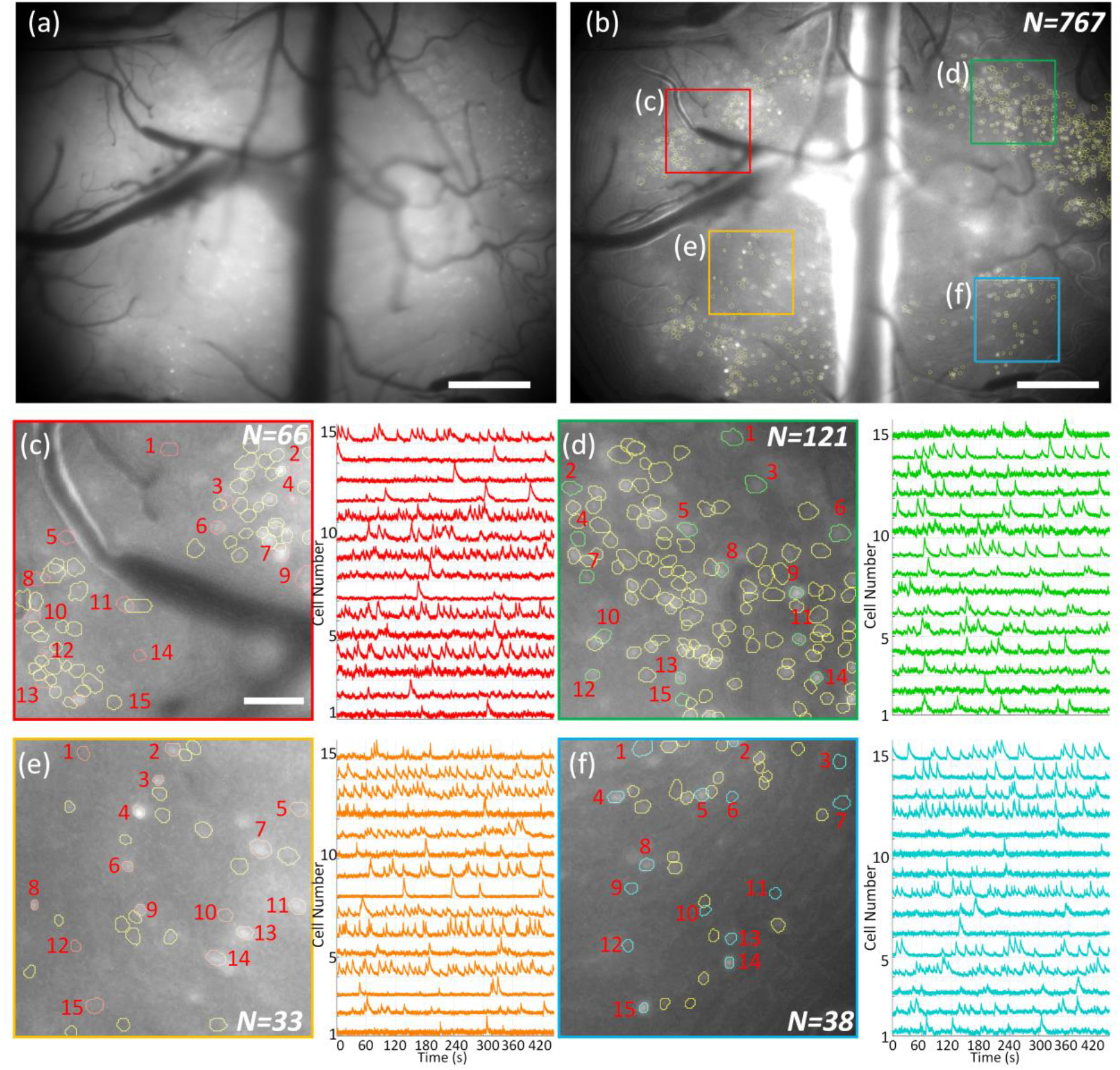
Ca^2+^ imaging of dorsal cortex through a midcranial window in an additional head-fixed mouse. (**a**) Maximum projection from a 8-minute recording session after motion correction. Scale bar: 500 μm. (**b**) Maximum-intensity projection image of the raw video after motion correction and background removed with contours of 767 extracted cells from CNMF-E analysis via CaImAn circled in yellow. The colored boxes indicate four sub-regions which are zoomed in and shown in **c-f**. Scale bar: 500 μm. (**c**) Map of 66 cells in the red-boxed area in **b** and 15 randomly chosen cells with their low-pass filtered Ca^2+^ transients for the numbered cells are shown in red. Scale bar, 100 μm. (**d**) Map of 121 cells in the green-boxed area in **b** and 15 randomly chosen cells with their low-pass filtered Ca^2+^ transients for the numbered cells are shown in green. (**e**) Map of 33 cells in the orange-boxed area in **b** and 15 randomly chosen cells with their low-pass filtered Ca^2+^ transients for the numbered cells are shown in red. (**f**) Map of 38 cells in the cyan-boxed area in b and 15 randomly chosen cells with their low-pass filtered Ca^2+^ transients for the numbered cells are shown in cyan.

**Table 1.**
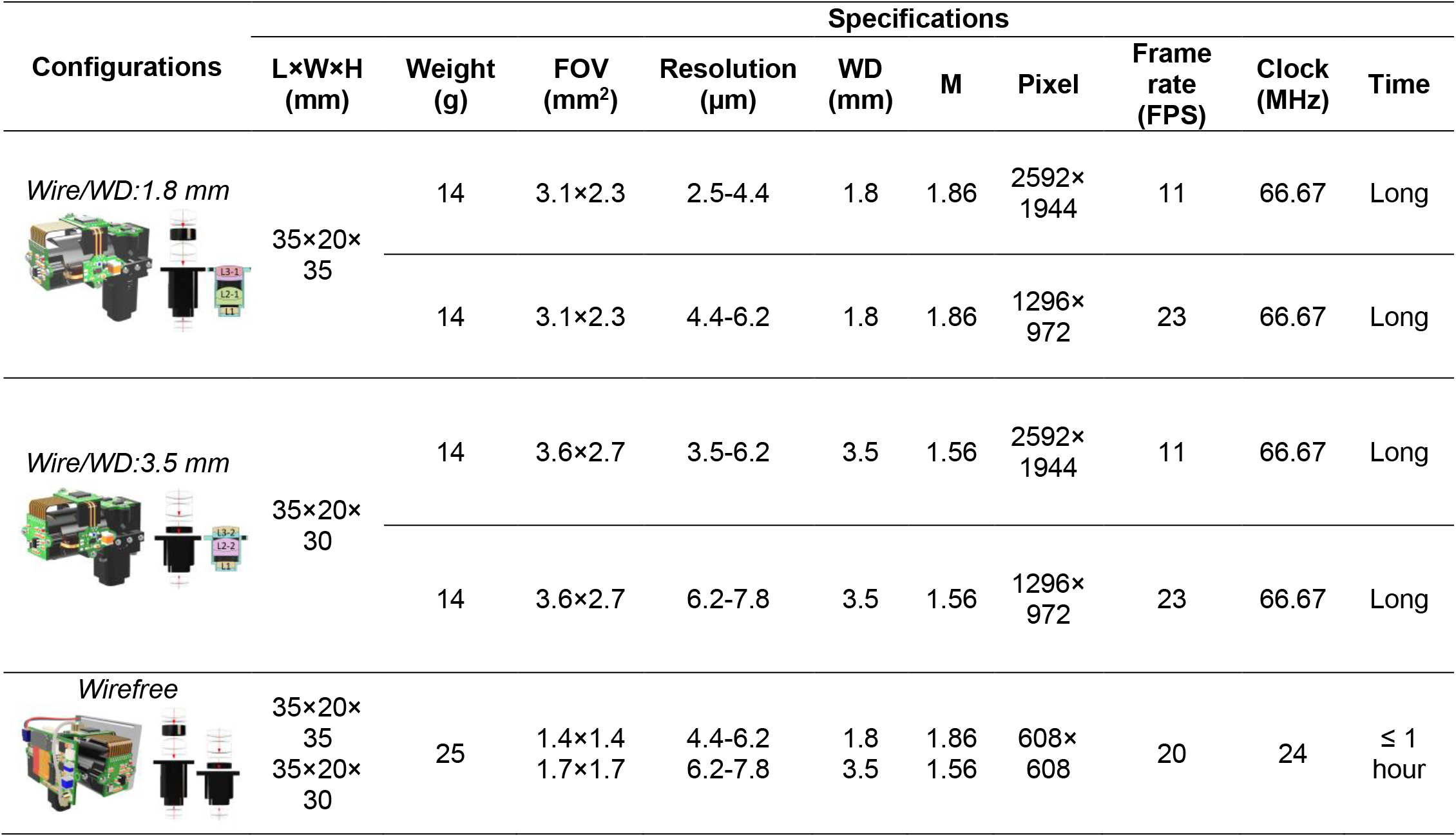
Three configurations of MiniLFOV.

| Configurations | Specifications |  |  |  |  |  |  |  |  |  |
| --- | --- | --- | --- | --- | --- | --- | --- | --- | --- | --- |
|  | LxWxH<br>(mm) | Weight<br>(g) | FOV<br>(mm<sup>2</sup>) | Resolution<br>(μm) | WD<br>(mm) | M | Pixel | Frame<br>rate<br>(FPS) | Clock<br>(MHz) | Time |
| *Wire/WD:1.8 mm* | 35x20x<br>35 | 14 | 3.1x2.3 | 2.5-4.4 | 1.8 | 1.86 | 2592x<br>1944 | 11 | 66.67 | Long |
|  |  | 14 | 3.1x2.3 | 4.4-6.2 | 1.8 | 1.86 | 1296x<br>972 | 23 | 66.67 | Long |
| *Wire/WD:3.5 mm* | 35x20x<br>30 | 14 | 3.6x2.7 | 3.5-6.2 | 3.5 | 1.56 | 2592x<br>1944 | 11 | 66.67 | Long |
|  |  | 14 | 3.6x2.7 | 6.2-7.8 | 3.5 | 1.56 | 1296x<br>972 | 23 | 66.67 | Long |
| *Wirefree* | 35x20x<br>35<br>35x20x<br>30 | 25 | 1.4x1.4<br>1.7x1.7 | 4.4-6.2<br>6.2-7.8 | 1.8<br>3.5 | 1.86<br>1.56 | 608x<br>608 | 20 | 24 | ≤ 1<br>hour |

**Table 2.**
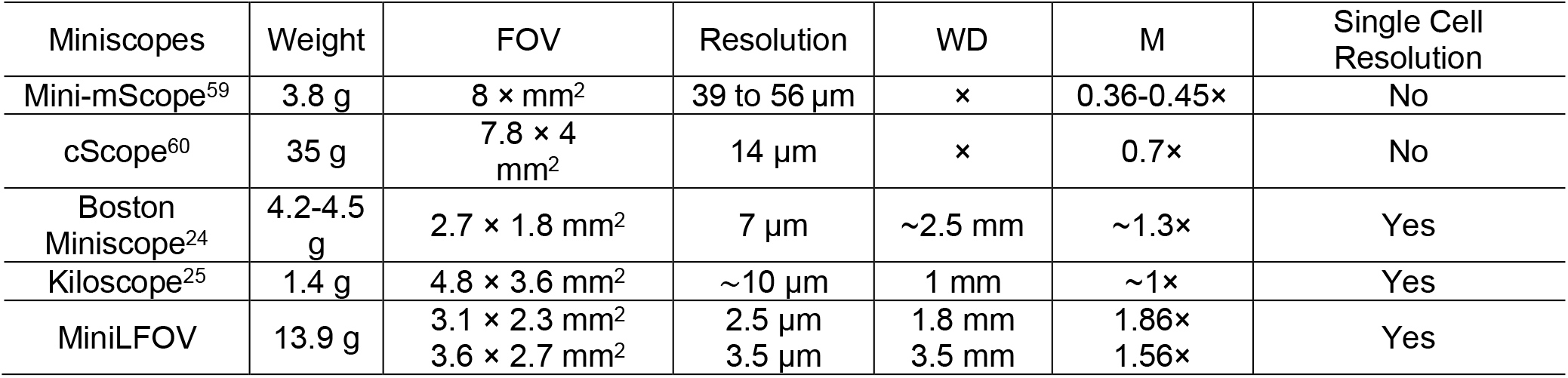
List of large FOV microscopes.

The goal of our proposed MiniLFOV is to develop a more stable, sensitive, and easy to access platform which can be broadly used in neuroscience research. Instead of prototype, proof-of-concept or custom design, MiniLFOV is designed with off-the-shelf lenses which are easy to get from Edmund optics. The weight is compromised to get a higher NA both for excitation and fluorescence collection to make it more sensitive than other large FOV counterparts. Higher sampling rate can be obtained with higher optical magnification.

**Table 3.**
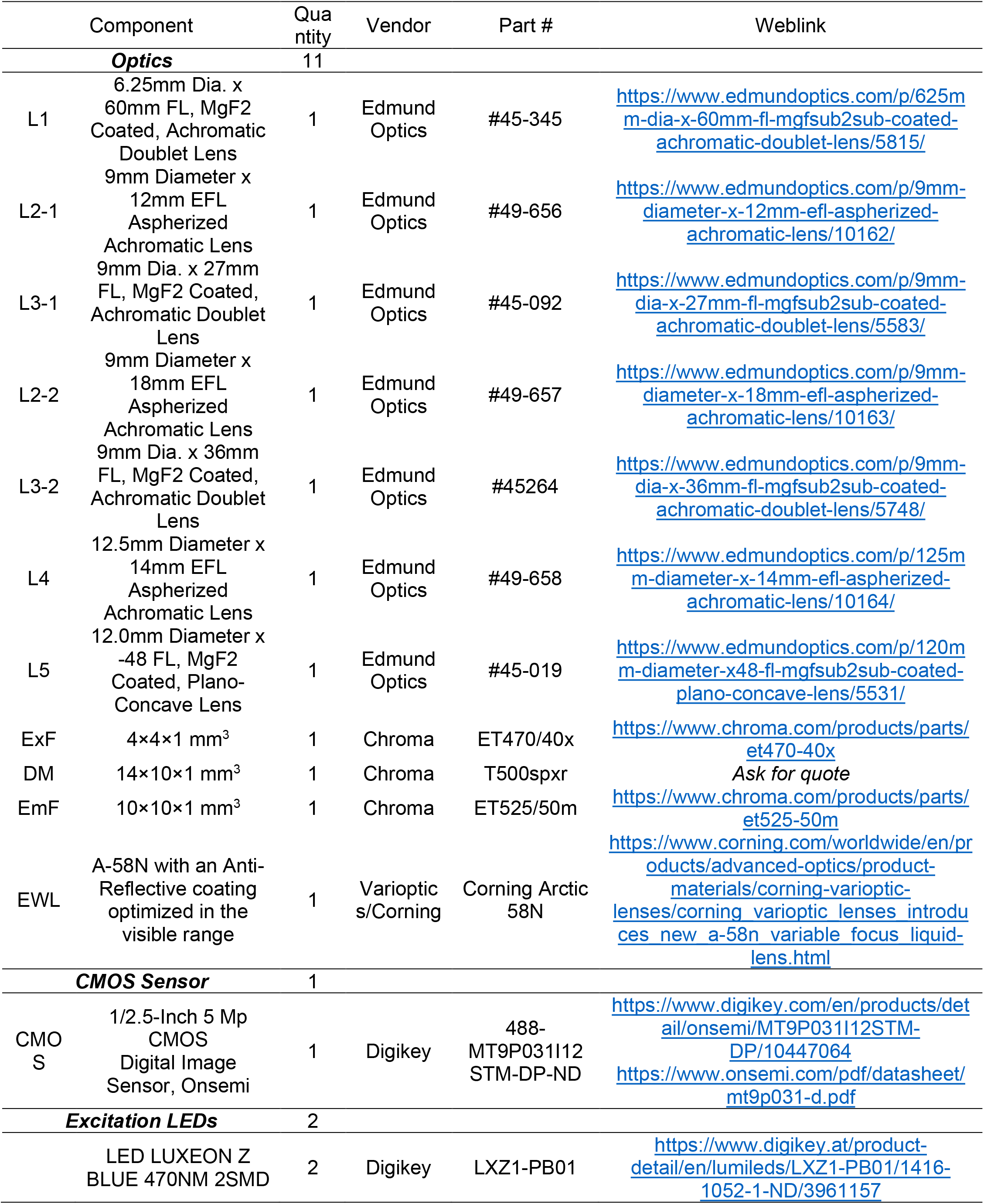
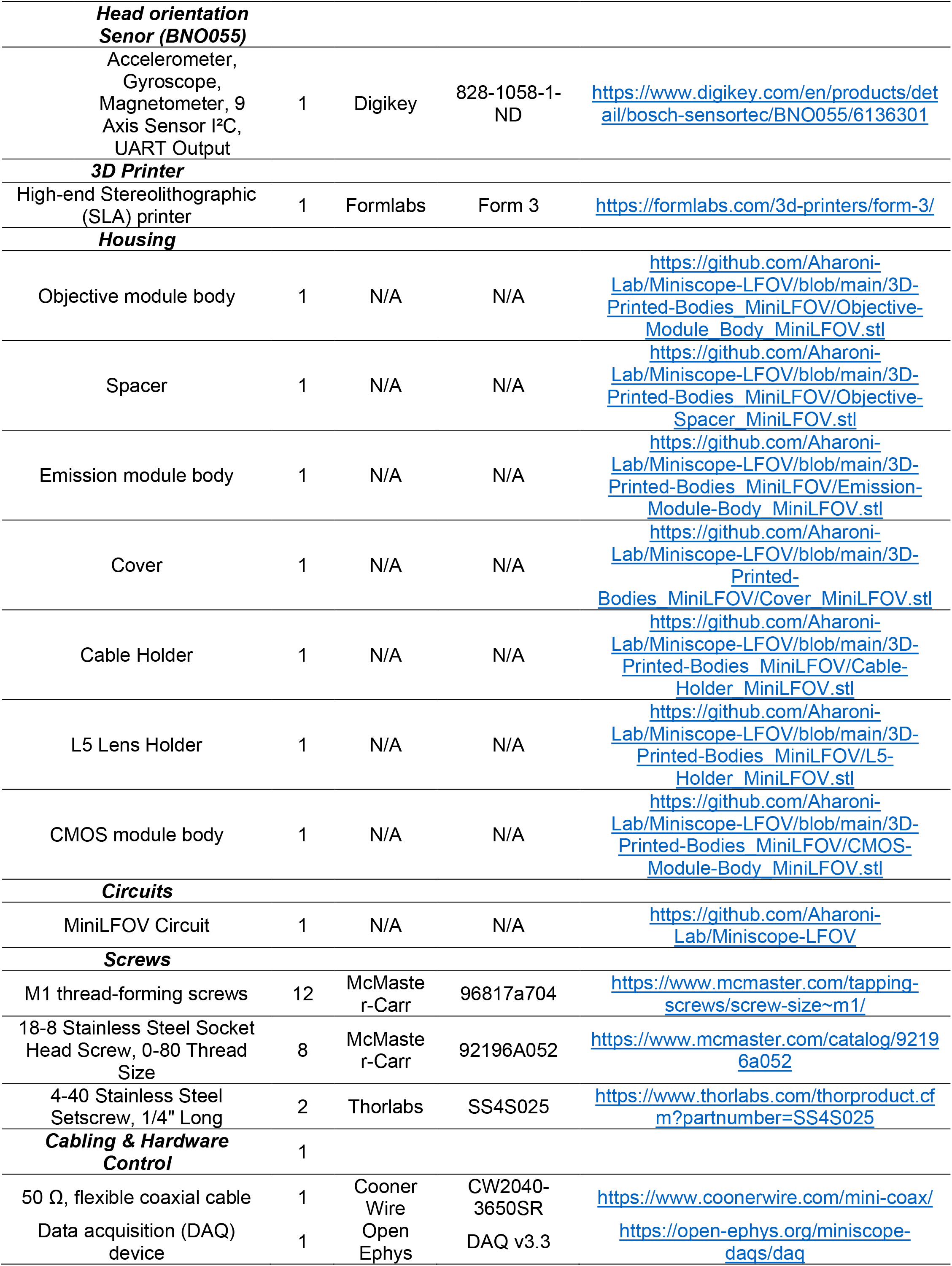
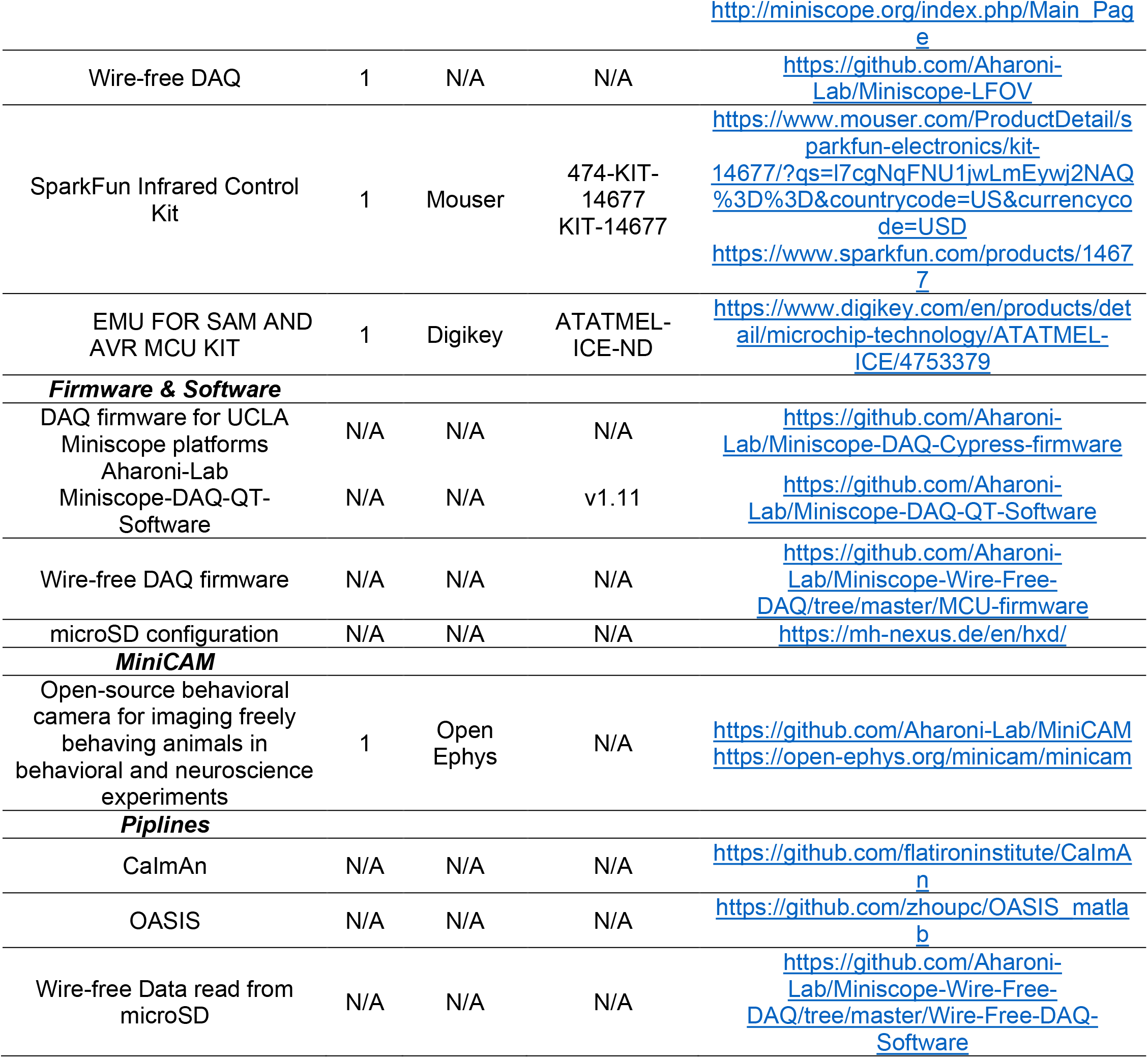
List of components for MiniLFOV assembly.

